# Changes in microbial composition, diversity, and functionality in the *Amblyomma maculatum* microbiome following infection with *Rickettsia parkeri*

**DOI:** 10.1101/2021.10.25.465777

**Authors:** Abdulsalam Adegoke, Deepak Kumar, Khemraj Budachetri, Shahid Karim

## Abstract

**Background:** Ticks are the primary vectors for emerging and resurging pathogens of public health significance worldwide. Examining tick bacterial composition, diversity, and functionality across developmental stages and tissues is necessary for designing new strategies to control ticks and prevent tick-borne diseases.

**Methods:** A high-throughput sequencing approach was used to determine the influence of blood meal and *Rickettsia parkeri* infection on changes in *Amblyomma maculatum* microbiome composition, diversity, and functionality across the developmental timeline and in different tissues. Quantitative insight into microbial ecology analysis allowed us to determine microbial population structure, composition, and diversity. A non-metric multidimensional scaling, the sparse correlations for compositional data (SparCC) module, and phylogenetic investigation of communities by reconstruction of unobserved states 2 (PICRUSt2) software were used in the assessment.

**Results:** The *Amblyomma maculatum* microbiome comprises ten bacterial genera present across tick life cycle stages. Among the top ten bacterial genera (the core tick microbiome), *Rickettsia, Francisella,* and *Candidatus Midichloria* are the key players, with positive interactions within each developmental stage and adult tick organ tested. The bacterial abundances, based on the number of operational taxonomic units (OTUs), increase with blood meal in each stage, helping bacterial floral growth. The growth in bacterial numbers is related to highly abundant energy metabolism orthologs with blood meal, according to functional analysis. Whereas *R. parkeri* had a positive correlation with *Candidatus Midichloria* during the tick life cycle, based on the increased number of OTUs and network analysis, this was due to an increased level of metabolic activity.

Interestingly, *R. parkeri* replaces *Francisella,* based on the lower level of OTUs representing *Francisella* in *R. parkeri*-infected ticks (in all stages/organs) and negatively correlated according to network and linear discriminant analysis effect size (LEfSe).

**Conclusions:** We found that *Rickettsia* and *Francisella* predominate in the core microbiome of the Gulf Coast tick, whereas *Candidatus Midichloria* and *Cutibacterium* levels increase with infection. Network analysis and functional annotation suggest that *R. parkeri* interacts positively with *Candidatus Midichloria* and negatively with *Francisella* and that metabolic profiles are upregulated with blood meal and *R. parkeri* infection. Overall, this is the first study to determine the combinatorial outcome of blood meal and pathogen interaction on microbiome composition over the developmental stages of *Am. maculatum*. This new study expands on our existing knowledge of the *Am. maculatum* microbiome and further highlights the need to investigate pathogen–symbiont interactions between *R. parkeri* and *Francisella* or *Candidatus Midichloria* to facilitate the development of strategies for controlling tick-transmitted diseases.

Graphical Abstract

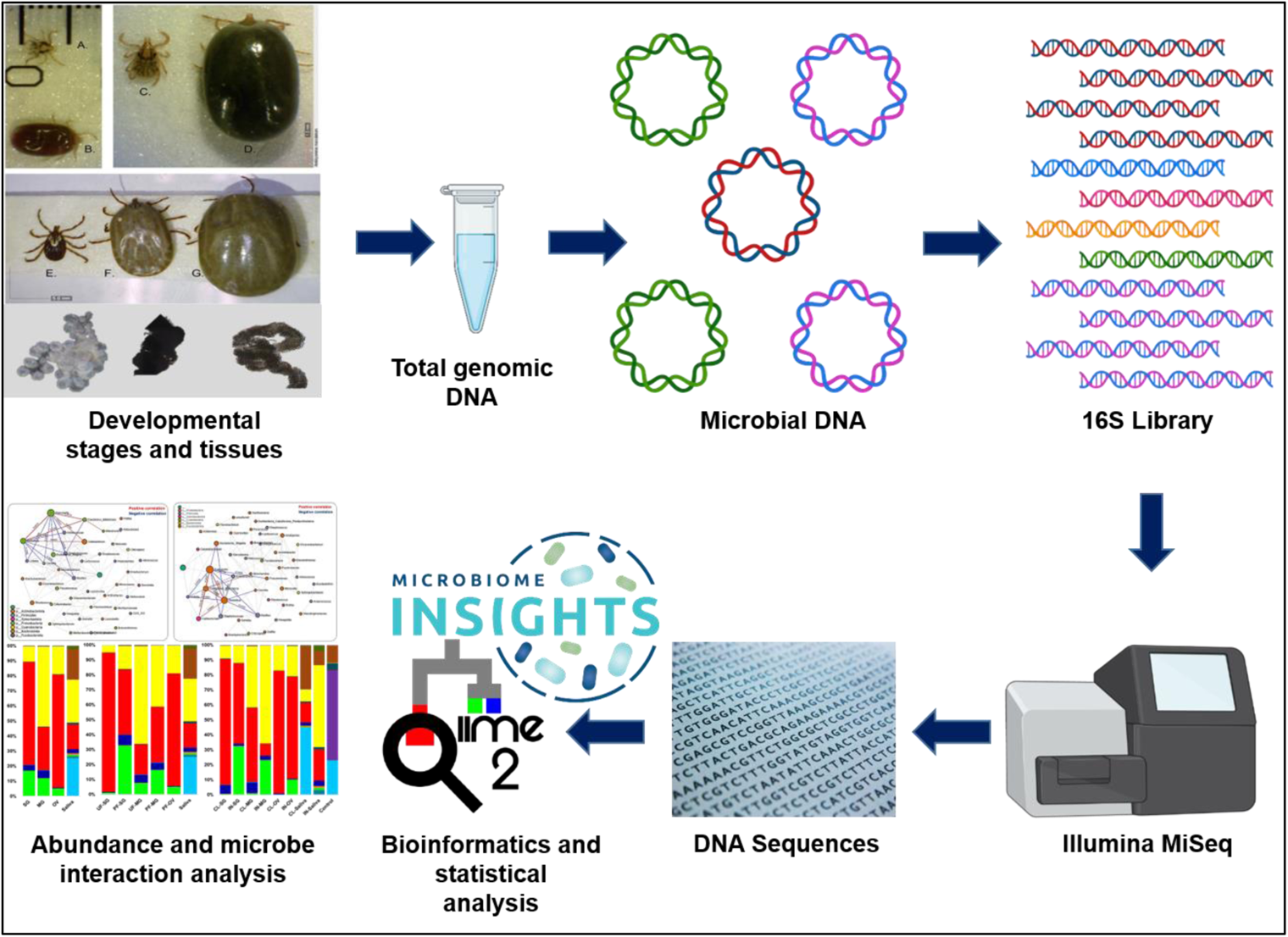

## Background

Ticks are obligate hematophagous arthropods that predominantly depend on human and animal host blood for development. Their importance as one of the important vectors of human and animal pathogens is highlighted by the substantial resources channeled towards controlling the several tick-transmitted pathogens. The Gulf Coast tick (*Amblyomma maculatum*) is one of four human-biting ticks in the United States, including also *Amblyomma americanum, Ixodes scapularis,* and *Dermacentor variabilis. Amblyomma maculatum* (*Am. maculatum*) is the most geographically widespread tick species in the southeastern United States (Sonenshine, 2018). It is the competent vector of *Rickettsia parkeri* (*R. parkeri*), a spotted fever group rickettsial (SFGR) pathogen that causes mild infections in the United States (Paddock and Goddard, 2015).

Recent advances in the control of tick and tick-transmitted pathogens have led to the discovery of microbial communities that co-exist alongside pathogens. The functional role of the tick microbiome has only been partly clarified by tick research. Overall, the primary focus of tick microbiome research has been on the microbial profiling of field-collected ticks to understand the basic bacterial profiles/communities present. The tick microbiome is a moving target and investigating its dynamics within the tick vector is a continuing concern (Narasimhan et al., 2015; Bonnet et al., 2017; Greay et al., 2018; Narasimhan et al., 2021). Ticks specialize in an exclusive diet of vertebrate blood and have evolved intimate interactions with beneficial symbionts that provide essential B vitamins and cofactors deficient in the blood diet (Guizzo et al., 2017; Ben-Yousaf et al., 2020; Duron et al., 2017; Bonnet et al., 2017; Duron et al., 2018; Gottlieb et al., 2015; Hunter et al., 2015; Olivieri et al., 2019; Smith et al., 2015), and the nutritional value of essential vitamins provided by the tick microbiome has been related to the reproductive fitness of ticks (Zhong et al., 2007). Immunomodulatory functions have also been associated with microbial communities within the tick and other blood-feeding arthropods, in which certain microbes have been shown to constantly prime the host immune system to continuously produce basal levels of antimicrobial peptides while also directly competing with or facilitating the establishment of different members of these microbial communities (de la Fuente et al., 2017; Fogaca et al., 2021; Boulanger and Wikel, 2021; Bonnet and Pollet, 2021; Budachetri et al., 2018).

Several studies have reported on the structure and microbial composition of field-collected and lab-maintained tick populations (Budachetri et al., 2014; Budachetri et al., 2017; Ross et al., 2018; Varela-Stokes et al., 2018; Thapa et al., 2019; Gill et al., 2020; Maldonado-Ruiz et al., 2021; Kumar et al., 2021; Karim et al., 2017; Zhang et al., 2019; Portillo et al., 2019; Adegoke et al., 2020; Guizzo et al., 2020; Rojas-Jaimes et al., 2021). These studies have revealed a variety of factors responsible for the differences among microbial communities between different tick samples, thus contributing extensively to our understanding of how microbial communities residing within the tick contribute to several aspects of tick biology (Ponnusamy et al., 2014; Budachetri et al., 2015; Trout-Fryxell and DeBruyn, 2016; Budachetri et al., 2018; Thapa et al., 2019; Brinkerhoff et al., 2020; Adegoke et al., 2020; Kueneman et al., 2021). However, the structure of *Am. maculatum* microbial communities and how these communities are shaped by blood meal and pathogen interactions have been the focus of only a few studies (Budachetri et al., 2014; Budachetri et al., 2018). Previous microbiome studies of *Am. maculatum* identified unique differences in microbial composition and abundance of specific bacterial genera in field-collected and questing tick tissues (Budachetri et al., 2014). The ability of a tick to serve as a competent vector for any pathogen goes beyond the establishment of the pathogen within the tick tissue but also extends to the facilitation of pathogen transmission at the tick–host interface, which is made possible in part by salivary proteins, in what has been described as saliva-assisted transmission (Nuttal and Labuda, 2008; Šimo et al., 2017; Nuttal, 2019; Karim et al., 2021). Whether the tick salivary microbiome interferes or facilitates pathogen colonization and transmission remains an unanswered question. While a study by Budachetri et al. (2014) reported several *Rickettsia* species in the saliva of *Am. maculatum*, only *R. parkeri* was detected in the salivary glands of *Am. maculatum*. Limited information exists about how microbial community members interact across tick developmental stages and tissues and how these tissues shape or modulate microbial community assemblages during tick physiological changes, such as those following blood-feeding or pathogen interactions. An elegant study by Lejal et al. (2021) identified several positive and negative correlations between members of the field-collected, questing *Ixodes ricinus* microbiome and how some of these interactions are modulated by tick-borne pathogens. It has been established that each tick tissue (organ) undergoes unique responses to specific physiological changes, but it is not known whether these responses also induce changes in the microbial communities residing within these tissues (Franta et al., 2010; Schwarz et al., 2014; Crispell et al., 2016; Kumar et al., 2016; Budachetri et al., 2017; Starck et al., 2018; Tirloni et al., 2020; Kurokawa et al., 2020). In the current study, a high-throughput sequencing approach was utilized to examine microbiome changes throughout the ontogeny of uninfected and pathogen-infected *Am. maculatum* ticks. Characterization of unique microbe-microbe interactions at the different tick developmental stages and tissues revealed potential synergistic and antagonistic interactions among the bacterial communities.

## Materials and Methods

### Ethics statement

All animal experiments were conducted in strict accordance with the recommendations of the Guide for the Care and Use of Laboratory Animals of the National Institutes of Health, USA. The protocols for the blood-feeding of immature and mature development stages of *Am. maculatum* was approved by the Institutional Animal Care and Use Committee of the University of Southern Mississippi (protocols #15101501.2 and 15011402).

### Tick maintenance

*Rickettsia parkeri*-infected and uninfected *A. maculatum* ticks were maintained under laboratory conditions as previously described (Budachetri et al., 2017; 2018). The colonies were established from field-collected adult ticks collected in 2011 from the Mississippi Sandhill Crane, National Wildlife Refuge (https://www.fws.gov/refuge/mississippi_sandhill_crane/) using the drag-cloth method. To generate replete adult females, unfed male and female *Am. maculatum* ticks were infested on sheep and allowed to feed to repletion and drop off. Fully engorged female ticks were then allowed to oviposit in individual snap-cap vials with perforated lids protected by breathable mesh cloths to allow for oxygen exchange while preventing their escape. Unfed larvae that emerged from the eggs were infested on golden Syrian hamsters until fully engorged, which were then allowed to molt into nymphs and then reinfested on golden Syrian hamsters. The fully fed nymphs were subsequently allowed to molt into adult male and female ticks. The presence or absence of *R. parkeri* at each developmental stage was confirmed following previously established nested PCR approaches (Budachetri 2014).

### Tick life stages and DNA extraction

DNA was isolated from freshly laid eggs as well as unfed and partially fed ticks at various developmental stages, including larvae, nymph, male, and female adults. To remove surface contaminants and bacterial carryover, all samples from developmental stages were surface-sterilized using 2% sodium hypochlorite, followed by two 5-minute washes in 70% ethanol and two 5-minute washes in sterile deionized water (Kumar et al., 2021). Tissues were dissected from unfed and partially blood-fed adult females and directly stored in ATL buffer before extracting DNA. Tick dissection was done in a biosafety cabinet while changing blades between dissections. Saliva was collected on ice by endogenous induction of salivation following injection of pilocarpine, as previously described (Ribeiro et al., 2004; Valenzuela et al., 2004). DNA was extracted using the DNeasy Blood and Tissue Kit following the manufacturer’s protocol (QIAGEN, Germantown, MD, USA) in a biosafety cabinet to eliminate potential carryover of bacteria between samples. Control samples included a blank control and a mock bacterial sample with a known bacterial population (ZymoBIOMICS™ Microbial Community DNA Standard) as part of the DNA-extraction protocol.

### Library preparation for Illumina 16S sequencing

Library preparation and indexing were carried out according to Illumina 16S metagenomic sequencing. The V3–V4 region of the 16S rRNA gene was amplified using DNA samples from different life stages and tissues. A two-step PCR amplification was carried out, which included an initial V3–V4 amplification using primers with specific Illumina adapter overhang nucleotide sequences using the following thermocycling conditions: 95°C for 3 min; followed by 25 cycles of 95°C for 30 s, 55°C for 30 s, and 72°C for 30 s; and a final extension step of 72 °C for 5 min. The PCR amplicon was briefly purified using AmPure XP beads (Agencourt Bioscience Corporation, Beverly, Massachusetts, USA) following the manufacturer’s protocol, and the purified product was eluted in 50 µl of TE buffer (Qiagen cat. no. 19086) and analyzed on a 2% gel to confirm a single DNA band with an estimated amplicon size of 550 bp. The second and most crucial step involved attaching unique indexes, in the form of nucleotide sequences, to the purified V3–V4 PCR product. Commercially available dual indexes (i5 and i7) from the Illumina Nextera index kit V2 (Illumina, San Diego, CA) were used for the second (index) PCR. Briefly, a reaction including each of the forward and reverse Nextera index primers, 1X KAPA HiFi HotStart ReadyMix (Kapa Biosystems, Wilmington, MA, USA), and 5 μl of each purified V3–V4 amplicon was set up using the following thermocycling conditions: 95°C for 3 min; followed by 8 cycles of 95°C for 30 s, 55°C for 30 s, and 72°C for 30 s; and a final extension step of 72 °C for 5 min. The second (index) PCR product was briefly purified using AmPure XP beads (Agencourt Bioscience Corporation, Beverly, Massachusetts, USA), following the manufacturer’s protocol. The purified product was eluted in 25 µl of TE buffer and analyzed on a 2% gel to confirm a single DNA band with an estimated amplicon size of 630 bp. The purified indexed PCR library products were quantified using the KAPA Library Quantification Kit from Roche (cat. no. 07960255001, kit code KK4844), normalized to a concentration of 7 nM, and pooled with 5 µl of each normalized library. Biological replicates from DNA extraction controls, TE buffer, and mock bacterial communities (ZymoBIOMICS™ Microbial Community DNA Standard) were simultaneously processed alongside the tick samples. The purified library was sequenced in a single run of an Illumina MiSeq sequencing instrument using reagent kit v2 (500 cycles) with 2 X 250 bp output at the University of Mississippi Medical Centre (UMMC) Genomics Core Facility.

### Data processing and analysis

Unless otherwise stated, all preprocessing was done following the video tutorial of the Quantitative Insights into Microbial Ecology (QIIME2) pipeline (Blyen et al., 2019). Briefly, demultiplexed fastq files were unzipped using the “unzip -q casava-18paired-end-demultiplexed.zip” command, followed by merging individual forward and reverse fastq files into a single fastq file. The Atacama soil microbiome pipeline was incorporated to control demultiplexed paired-end reads using the DADA2 plugin as previously described (Callahan et al., 2016). Low-quality sequences were trimmed and filtered out, and subsequent merging of paired-end reads ensured 20 nucleotide overhangs between forward and reverse reads. Chimeric sequences were removed from the sequence table. The qiime2 feature-classifier plugin v. 7.0 was used for taxonomic assignment against the pre-trained Silva_132_99% classifier (Quast et al., 2013). The “qiime diversity core-metrics-phylogenetic” command was used to compute diversity metrics, after which the “QIIME diversity alpha-group-significance” command was used to explore microbial diversity using Faith_pd and evenness metrics as measures of community richness and evenness, respectively. Raw sequences will be submitted to the required databases.

### Functional characterization of microbial communities

To determine the functionality of the microbial genome, we predicted the functional pathways using the Phylogenetic Investigation of Communities by Reconstruction of Unobserved States 2 (PICRUSt2) pipeline. We assigned functional predictions by normalizing the bacterial sequences and OTUs to the KEGG Ortholog (KO) databases and Clusters of Orthologous Groups (COGs) to predict metabolic pathways.

### Visualization

Microbiome Analyst, a web-based interface, was used for data visualization using taxonomy and metadata tables generated from data processing as input files (Dhariwal et al., 2017; Chong et al., 2020). Low count and prevalence data were filtered from the OTU table by setting values of 10 and 20, respectively. A filtered abundance table was exported and used in generating histograms of bacterial abundance in Microsoft Excel 2016 (Microsoft, 2018). Boxplots of alpha diversity were generated with GraphPad Prism version 8 for Windows (GraphPad Software, La Jolla California USA, www.graphpad.com) using the pairwise table generated from the “qiime diversity alpha-group-significance” command. Network correlation maps were constructed based on the sparse correlations for compositional data (SparCC) approach (Friedman and Alm, 2012). This approach uses the log-transformed values to carry out multiple iterations, which subsequently identifies taxa outliers to the correlation parameters (Chong et al., 2020). To compare the differences in the microbiome between tick groups based on measures of distance or dissimilarity, a matrix was generated from log-transformed sequence data, and ordination of the plots was visualized using both Principal Coordinates Analysis (PCoA) and non-metric multidimensional scaling (NMDS). The Bray–Curtis distance matrix was used to visualize compositional differences in the microbiome across all groups.

### Statistical Analysis

The Wilcoxon rank-sum test estimated significant correlations in diversity indexes across developmental stages and tissue data sets. A Kruskal–Wallis test followed by Dunn’s multiple comparison test was used in comparing the effect of blood meal and *R. parkeri* on microbial richness between two different groups (unfed vs. fed or uninfected vs. infected).

## Results

### Sequencing results

A total of 91 samples, representing at least three biological replicates across developmental stages (eggs, larvae, nymph, and adult), tissues (salivary gland, midgut, and ovaries), and body fluids (saliva) from uninfected and *R. parkeri*-infected ticks, generated a total of 4,090,811 forward and reverse reads, with a mean of 45,453.5 and a maximum of 112,242 reads per sample (Additional File, Table 1). Rarefaction analysis of the individual samples from a sequencing depth of 500 to 4000 confirmed that there was adequate sequence coverage relative to the number of operational taxonomic units (OTUs) and a plateau for individual tick samples (Additional File, Fig. 1A, B). Overall, 711 unique OTUs were identified, with 254 identified from each developmental stage and tissue.

**Figure 1:**
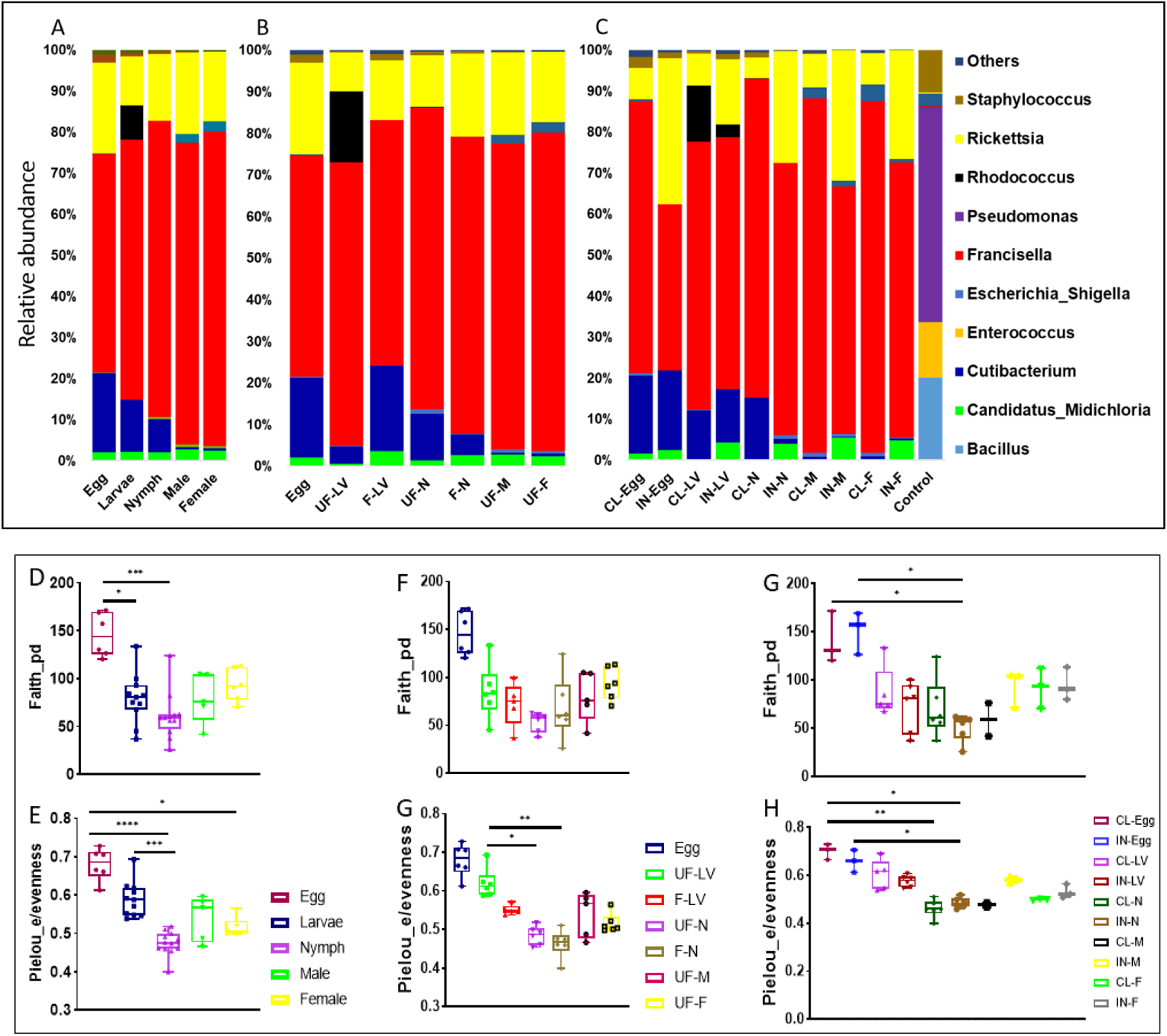
Stability of bacterial abundances and microbial richness across developmental stages. The relative abundances of the predominant members of microbial communities in A) all developmental stages, B) unfed and fully fed developmental stages and C) *R. parkeri*-infected and uninfected developmental stages. Diversity analysis using Faith phylogenetic distance (species richness) and Pielou_e metrics (evenness) in (D, E) all developmental stages, (F, G) unfed and fully fed developmental stages, and (H, I) *R. parkeri*-infected and uninfected developmental stages. Each bar on the abundance plot represents average data from 3–5 individual replicates. Each point of the box plots represents average data from 3–5 individual replicates. Unfed (UF), fed (F), larvae (LV), nymph (N), male (M), female (F), clean (CL), infected (IN).

### Microbiome dynamics across developmental stages

Proteobacteria represented the most abundant phylum throughout ontogeny, with a relative abundance of 84%, followed by Actinobacteria, Firmicutes, and an uncharacterized phylum (Additional File, Fig. 2A). An extended breakdown of each identified phylum and the number of member bacteria at the genus level is represented using pie charts for each phylum, as shown in Additional File, Fig. 2B–D. The breakdown of the top 10 most frequent bacterial genera shows that *Francisella* and *Rickettsia* were the predominant members of the *Am. maculatum* microbiome across tick developmental stages (Fig. 1A). We observed a progressive decrease in the relative abundance of the genus *Cutibacterium* throughout each developmental stage, from egg to adult male and female ticks (Fig. 1A). Other bacterial genera, such as *Rhodococcus*, *Pseudomonas*, and *Candidatus*_*Midichloria*, were also detected at relatively lower abundance. The abundance of representative bacterial genera was also compared between blood-fed and unfed life stages. Uptake of a blood meal was observed to impact the relative abundance of *Francisella* and *Cutibacterium* (Fig. 1B). Following a blood meal, the abundance of *Francisella* was negatively impacted, whereas *Cutibacterium* abundance went up in blood-fed larvae compared with unfed larvae.

**Figure 2:**
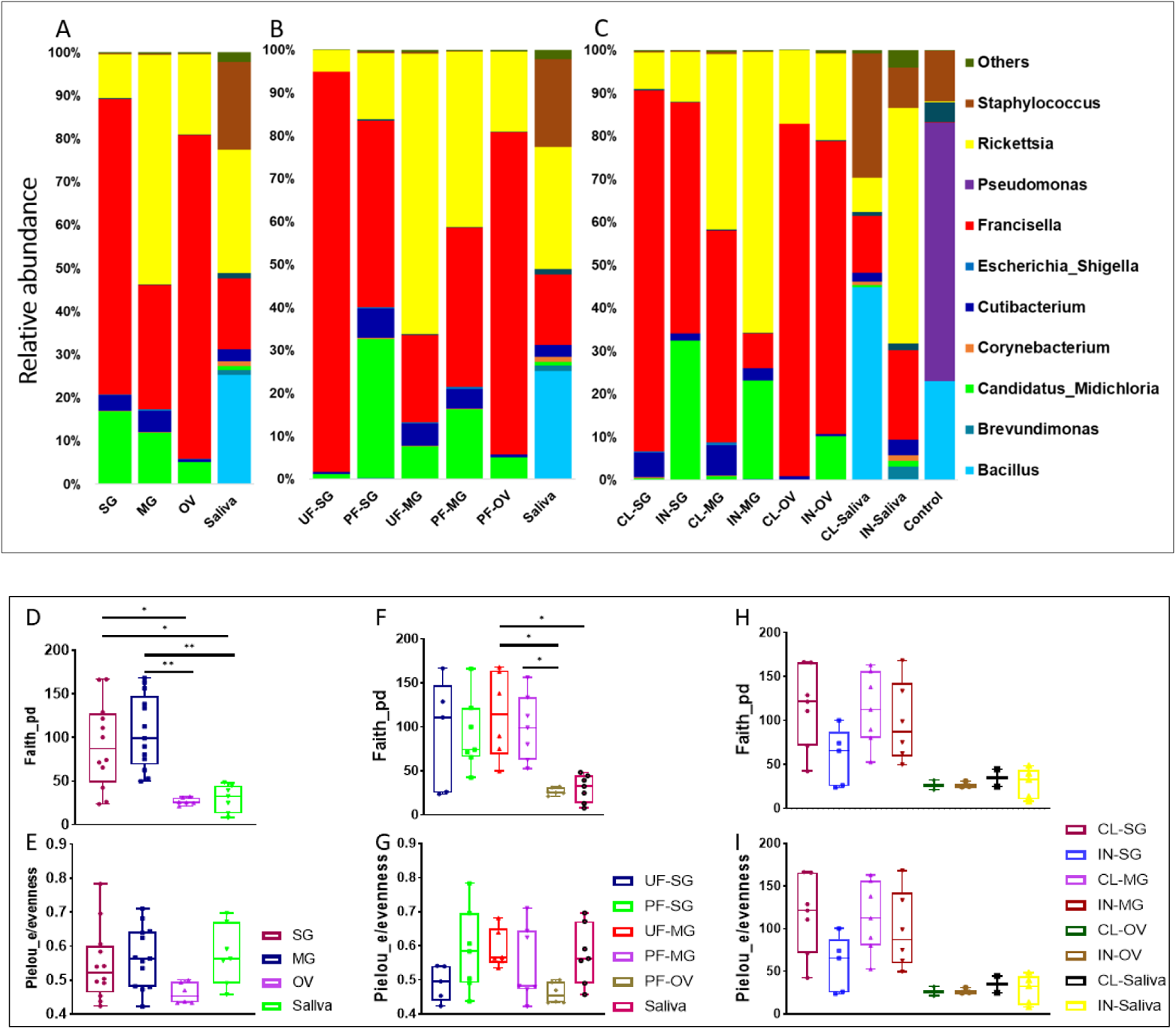
Changes in microbial assemblages and diversity across different tissues. The relative abundances of the predominant members of microbial communities in A) all isolated tissues, B) unfed and fully fed tissues, and C) *R. parkeri*-infected and uninfected tissues. Diversity analysis using Faith phylogenetic distance (specie richness) and Pielou_e metrics (evenness) in (D, E) all tissues, (F, G) unfed and fully fed tissues, and (H, I) *R. parkeri*-infected and uninfected tissues. Each bar on the abundance plot represents average data from 3–5 individual replicates. Each point of the box plots represents average data from 3–5 individual replicates. Unfed (UF), fed (F), larvae (LV), nymph (N), male (M), female (F), clean (CL), infected (IN). salivary gland (SG), midgut (MG), ovary (OV).

Interestingly, opposing trends were observed when blood-fed nymphs were compared with unfed nymphal ticks. The relative abundance of *Cutibacterium* was reduced, whereas *Francisella* abundance was observed to increase (Fig. 1B). To examine the modulation of microbial assemblages in the presence of pathogenic microbes, we compared the relative abundances of bacteria between uninfected and *R. parkeri*-infected samples from each developmental stage. There is a clear trend of decreasing *Francisella* abundance, and a corresponding increase in *Rickettsia* and *Candidatus*_*Midichloria* across all *R. parkeri*-infected stages compared with uninfected samples (Fig. 1C). Notably, the *Cutibacterium* load was negatively impacted by the presence of *R. parkeri* when infected and uninfected larvae and nymphs were compared (Fig. 1C).

Eggs exhibited higher bacterial richness and evenness, regardless of blood meal or infection (Wilcoxon rank-sum test *p < 0.05*). At the same time, nymphs showed the least microbial richness and evenness (Wilcoxon rank-sum test *p < 0.001*) (Fig. 1D, E). No significant microbial richness in the blood-fed samples was observed; however, blood meal was seen to exert a significant impact on species proportions in unfed nymphs when compared with unfed larvae (Kruskal–Wallis test followed by Dunn’s multiple comparison test, *p < 0.05*) and in the fed nymphs compared with unfed larvae (Kruskal–Wallis test followed by Dunn’s multiple comparison test, *p < 0.001*) when the Pielou_e evenness index was estimated (Fig. 1F, G) Overall, blood-feeding had no significant effect on microbial richness compared with unfed ticks across developmental stages.

The presence of *R. parkeri* infection was not found to affect microbial richness and evenness between developmental stages. However, a reduction in microbial richness in *R. parkeri*-infected nymphs compared with uninfected eggs (Kruskal–Wallis test followed by Dunn’s multiple comparison test, *p < 0.05*) and infected eggs (Kruskal–Wallis test followed by Dunn’s multiple comparison test, *p < 0.05*) was noted (Fig. 1H). *R. parkeri* infection was seen to reduce species proportion when an uninfected nymphal sample was compared with uninfected egg samples (Kruskal–Wallis test followed by Dunn’s multiple comparison test, *p < 0.001*); likewise, when the infected nymphal sample was compared with an uninfected egg sample (Kruskal–Wallis test followed by Dunn’s multiple comparison test, *p < 0.05*) and when an uninfected nymphal sample was compared with an infected egg sample (Kruskal–Wallis test followed by Dunn’s multiple comparison test, *p < 0.05*) (Figure 1I). NMDS analysis of community structure using the Bray–Curtis (BC) distance matrix (PERMANOVA, F-value = 22.818; R^2^ = 0.75015; p-value < 0.001 [NMDS] Stress = 0.12544) revealed two distinct clustering patterns in which the earlier developmental stages (egg, larvae, and a few nymphs) uniquely clustered away from the later developmental stages (male, female, and a few nymphs) (Additional File, Fig. 3). The BC distance matrix measures dissimilarity in microbial communities based on compositional differences among our samples. The observed clustering pattern further supports the differences observed in the presence of *Cutibacterium* in the eggs, larvae, and nymphs but not in adult male and female ticks. Overall, both *Francisella* and *Rickettsia* predominated in the microbiome during the developmental stages. Both blood meal and *R. parkeri* infection negatively affected the abundance of *Francisella,* while blood meal reduced microbial richness in earlier developmental stages.

**Figure 3:**
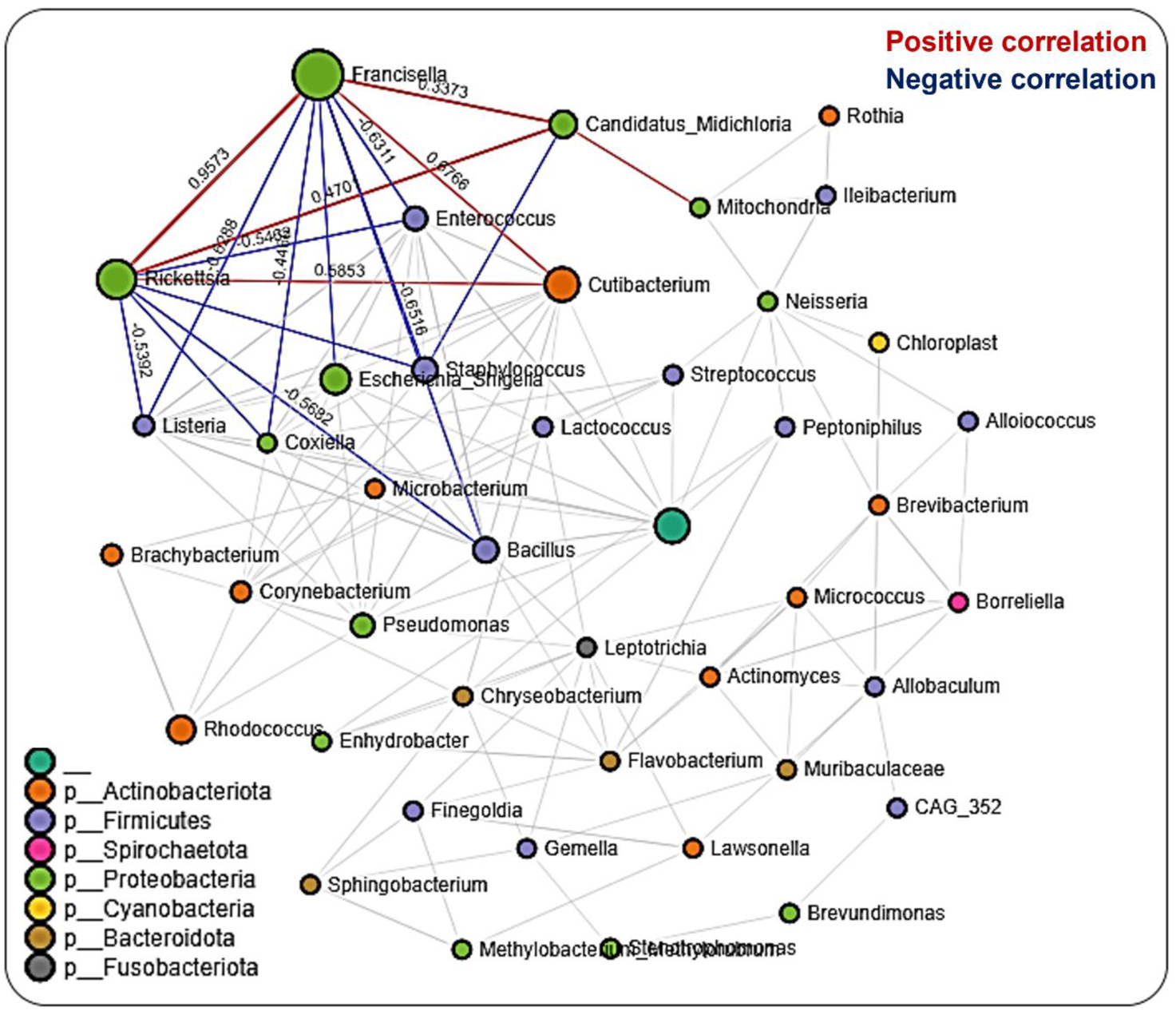
Correlation network analysis across the life stages of *Am. maculatum* ticks. Correlation network generated using the SparCC algorithm. Correlation network with nodes representing taxa at the family level and edges representing correlations between taxa pairs. Node size correlates with the number of interactions in which a taxon is involved, and the line color represents either positive (red) or negative (blue) correlations. The color-coded legend shows the bacterial family to which each taxon belongs.

### Microbiome composition and diversity across tissue niches

As with developmental stages, the phylum Proteobacteria had the greatest abundance of bacteria, with a relative abundance of 86%, followed by Firmicutes, Actinobacteria, and an unknown phylum, as illustrated in the Additional File, Fig. 3A–D. Representative sequences generated from tissue samples showed a similar trend to what was found for developmental stages. *Francisella* and *Rickettsia* predominated in the microbiomes of isolated tissues. In addition, the genus *Candidatus*_*Midichloria* was detected at relatively high abundance across isolated tissues, except for the saliva microbiome, which contained the genus *Bacillus*. By contrast, the genus *Cutibacterium* was present in all tissues, except for ovarian samples (Fig. 2A). Following blood uptake, the proportion of bacteria identified in the partially fed salivary gland was higher than in the unfed salivary gland (Fig. 2B). Interestingly, blood meal reduced the abundance of *Francisella*, whereas the prevalence of *Rickettsia, Cutibacterium,* and *Candidatus*_*Midichloria* increased (Fig. 2B). The effect of blood meal on the microbial abundance in the midgut tissues was different from salivary gland; in the reverse of what was observed in the salivary gland, blood meal increased the relative abundance of *Francisella* and *Candidatus*_*Midichloria*, while reducing that of *Rickettsia*. Due to the unique developmental trajectory of the female tick, in which ovarian development did not commence until 4–5 days post-blood-feeding, ovarian tissues from unfed female ticks were not included in this analysis (Fig. 2B). Bacterial communities were significantly different between unfed and partially blood-fed salivary gland (PERMANOVA F-value = 22.766; R^2^ = 0.7779; p-value < 0.001) and midgut tissues (PERMANOVA F-value = 16.167; R^2^ = 0.69785; p-value < 0.001), according to PCoA analysis of the Bray–Curtis distance matrix (Additional File, Fig. 4A, B).

**Figure 4:**
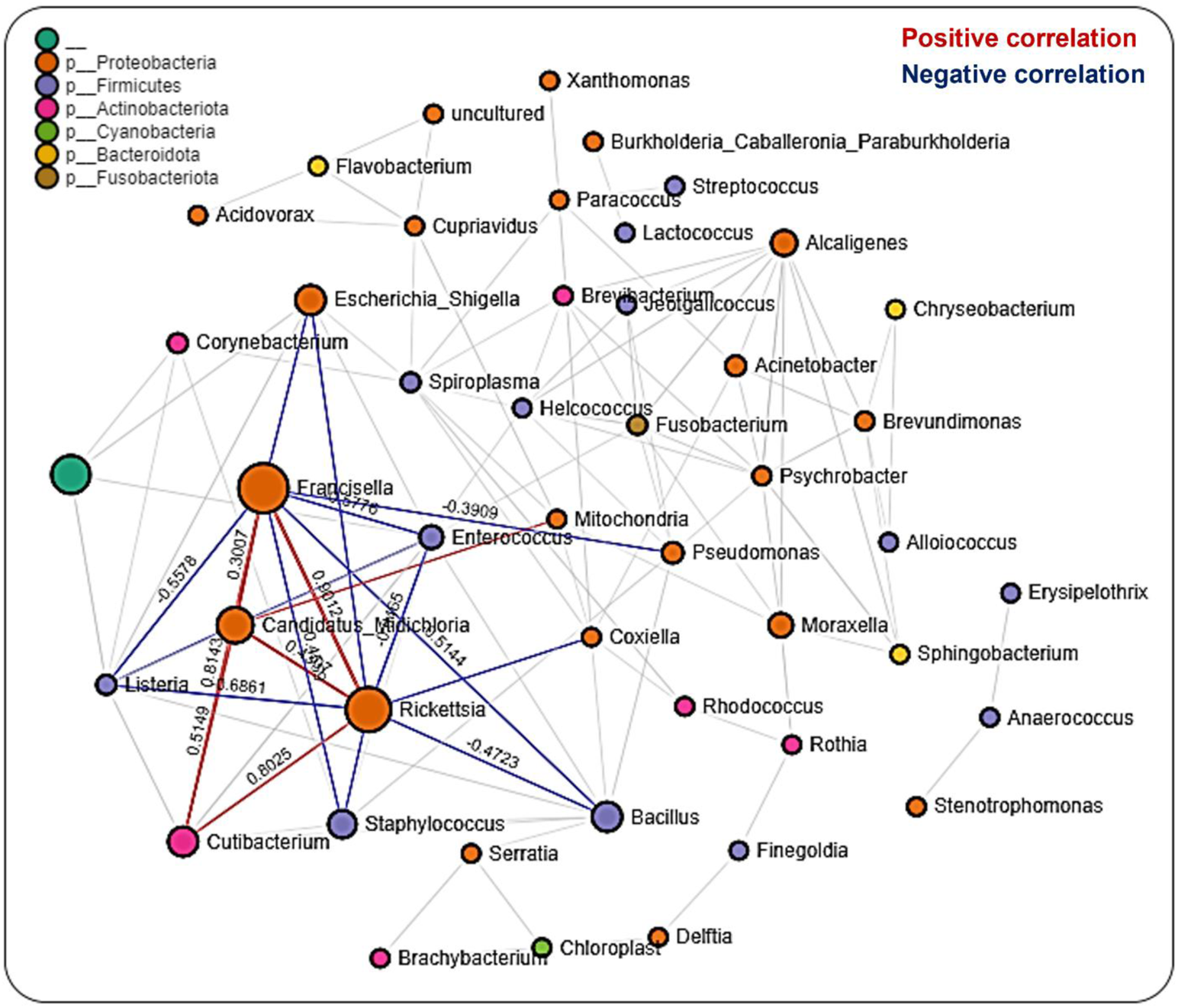
Correlation network analysis across tissue stages of *Am. maculatum* ticks. Correlation network generated using the SparCC algorithm. Correlation network with nodes representing taxa at the family level and edges representing correlations between taxa pairs. Node size correlates with the number of interactions in which a taxon is involved, and the line color represents either positive (red) or negative (blue) correlations. The color-coded legend shows the bacterial family to which each taxon belongs.

The presence of *R. parkeri* was associated with a corresponding increase in the relative abundance of *Candidatus*_*Midichloria* in all infected compared with uninfected tissues, with a corresponding decrease in the prevalence of *Francisella* (Fig. 2C). There was an increase in the relative abundance of *Rickettsia* detected in *R. parkeri*-infected saliva compared with saliva from uninfected ticks. Bacterial communities were significantly different between uninfected and infected salivary gland (PERMANOVA F-value = 20.316; R^2^ = 0.75761; p-value < 0.001), midgut (PERMANOVA] F-value = 36.225; R^2^ = 0.83806; p-value < 0.001) and ovarian tissues (PERMANOVA F-value = 29.509; R^2^ = 0.89397; p-value < 0.008), according to PCoA analysis of the Bray–Curtis distance matrix (Additional File, Fig. 5A–C).

**Figure 5:**
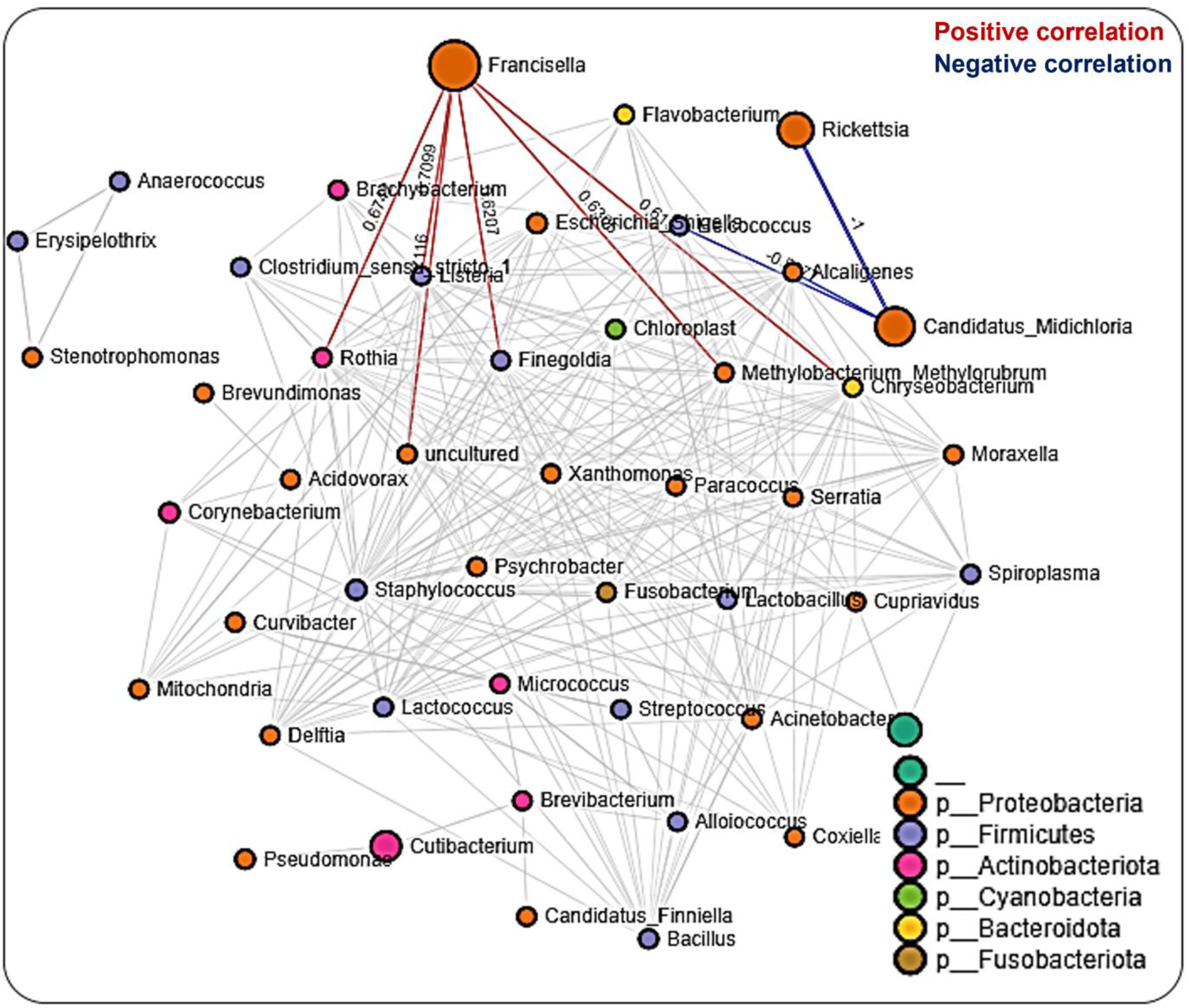
Correlation network analysis in the salivary glands of *Am. maculatum* ticks. Correlation network generated using the SparCC algorithm. Correlation network with nodes representing taxa at the family level and edges representing correlations between taxa pairs. Node size correlates with the number of interactions in which a taxon is involved, and the line color represents either positive (red) or negative (blue) correlations. The color-coded legend shows the bacterial family to which each taxon belongs.

Overall, microbial richness was highest in midgut tissues (Wilcoxon rank-sum test *p < 0.001*) and lowest in the ovary (Wilcoxon rank-sum test *p < 0.05*), based on the Faith_phylogenetic distance index. At the same time, no significant differences were found in microbial proportions across different tissues, including saliva (Fig. 2D, E). The impact of blood meal on alpha diversity between unfed and partially fed tissues was assessed, and blood meal was found to reduce the species richness following blood uptake in the midgut compared with the unfed midgut (Fig. 2F), while species proportions were not significantly impacted (Fig. 2G). Similarly, there was no significant difference in either species richness or evenness between *R. parkeri*-infected and uninfected tissues (Fig. 2H, I). Here we found that blood meal and *R. parkeri* infection shaped the *Am. maculatum* microbiome in a tissue-dependent manner.

### Microbiota interactions across developmental stages

We performed network analysis to visualize significant positive and negative correlations between the bacterial genera identified. We observed a total of 286 significant partial correlations between 43 bacterial genera, 52% (150) of which showed positive correlations, while the remaining 48% (136) exhibited negative correlations (Additional File, Table 2). All identified bacteria were from the phyla Actinobacteria (10 genera), Firmicutes (12 genera), Spirochaetota (1 genus), Proteobacteria (12 genera), Cyanobacteria (1 genus), Bacteroidota (4 genera), and Fusobacteria (1 genus) as well as an unknown phylum (1 genus) (Fig. 3). The genera *Francisella, Rickettsia,* and *Candidatus*_*Midichloria* were found within the same network cluster where they share significant positive correlations. Individually, they both interacted with other bacteria genera as is the case with *Francisella* and *Rickettsia* OTUs sharing negative correlations with *Listeria, Coxiella, Bacillus, and Enterococcus,* and positive correlations with Cutibacterium. The genus *Candidatus*_*Midichloria* on the other hand was positively correlated with OTUs belonging to the *Mitochondria* genus. It is worth noting that *Candidatus*_*Midichloria* had fewer taxa with which it interacted than *Francisella or Rickettsia* (Additional File, Table 2). Another interesting observation made from the network interactions was the significant negative correlations in the interactions between *Coxiella* and *Francisella* and between *Coxiella* and *Rickettsia*. A more surprising correlation was identified between *Bacillus, Listeria, Enterococcus,* and *Staphylococcus*, all of which are members of the phylum Firmicutes. They were all negatively correlated with *Rickettsia*, *Francisella,* and *Candidatus*_*Midichloria* from Proteobacteria (Fig. 3). To identify the significantly represented microbial genera between different life stages, we used linear discriminant analysis effect size (LEfSe) to determine the respective LDA scores for the identified bacteria observed in the samples. LDA analysis showed that, out of 46 bacteria taxa, 13 were significantly correlated at different life stages. Several of the bacteria either increased or decreased in more than one life stage. By contrast, the bacterial taxon *Enhydrobacter* is the only genus represented exclusively at the egg stage (Additional File, Table 3). Likewise, the taxa *Enterococcus, Lactobacillus,* and *Coxiella* were not detected at any developmental stages. Another interesting observation was the increased presence of the genera *Corynebacterium* and *Cutibacterium* exclusively at early developmental stages (Additional File, Fig. 4).

### Tissue-driven microbial interactions

Network analysis in tissues showed 220 significant correlations between 46 bacterial genera, each belonging to the phyla Proteobacteria (20 genera), Firmicutes (13), Actinobacteriota (6 genera), Cyanobacteria (2 genera), Bacteroidota (3 genera), and Fusobacteriota (1 genus) as well as an unknown phylum (1 genus). Seventy-two (33%) were negative correlations, while 148 (67%) were positive correlations (Additional File, Table 4). We observed that OTUs belonging to *Candidatus_Midichloria, Francisella* and *Rickettsia* were positively correlated and represent a network cluster that shares both positive and negative correlations with OTUs from other bacteria genera. Interestingly, bacteria belonging to the phylum Firmicutes, such as *Listeria, Staphylococcus, Bacillus, and Enterococcus* were negatively correlated with *Francisella, Rickettsia,* and *Candidatus*_*Midichloria* which are bacteria from the Proteobacteria phylum (Fig. 4). LDA analysis revealed that *Francisella* was significantly enriched in ovaries and salivary glands, whereas *Rickettsia* was increased significantly in the midgut tissues and saliva (Additional File, Table 5; Fig. 5). The saliva microbiome was also the only sample exclusively enriched with *Alcaligenes, Moraxella, Pseudomonas,* and *Spiroplasma*. The first three belong to the phylum Proteobacteria and the latter from the phylum Firmicutes.

### Microbial interactions in the salivary gland

A total of 608 significant partial interactions between 48 bacteria genera were identified in network analysis of the salivary gland, with 176 (29%) negative correlations and 432 (71%) positive correlations (Additional File, Table 6). Most interactions were seen with *Francisella, Rickettsia,* and *Candidatus*_*Midichloria* (Fig. 5). The most surprising correlation was the negative correlation observed between *Rickettsia* and *Candidatus*_*Midichloria*, and the most interesting was that *Rickettsia* shares a correlation network only with *Candidatus*_*Midichloria* (Fig. 5). No direct interaction was observed between *Francisella* and *Candidatus*_*Midichloria* or between *Francisella* and *Rickettsia*. LDA analysis of significantly enriched bacteria revealed a unique pattern between *R. parkeri*-infected and uninfected salivary glands. Uninfected salivary glands were exclusively enriched with *Francisella, Bacillus, Cutibacterium, Pseudomonas,* and *Stenotrophomonas.* By contrast, the genera *Enterococcus, Rickettsia,* and *Candidatus*_*Midichloria* were the only bacteria significantly enriched in *R. parkeri-*infected salivary glands (Additional File, Fig. 6A). Interestingly, *Francisella* was the only bacterium identified as significantly enriched in the unfed salivary gland. While *Francisella* was also enriched in the partially fed salivary gland, several other bacteria were identified, such as *Escherichia_Shigella, Bacillus, Staphylococcus, Enterococcus, Cutibacterium,* and *Pseudomonas* (Additional File, Fig. 6B).

**Figure 6:**
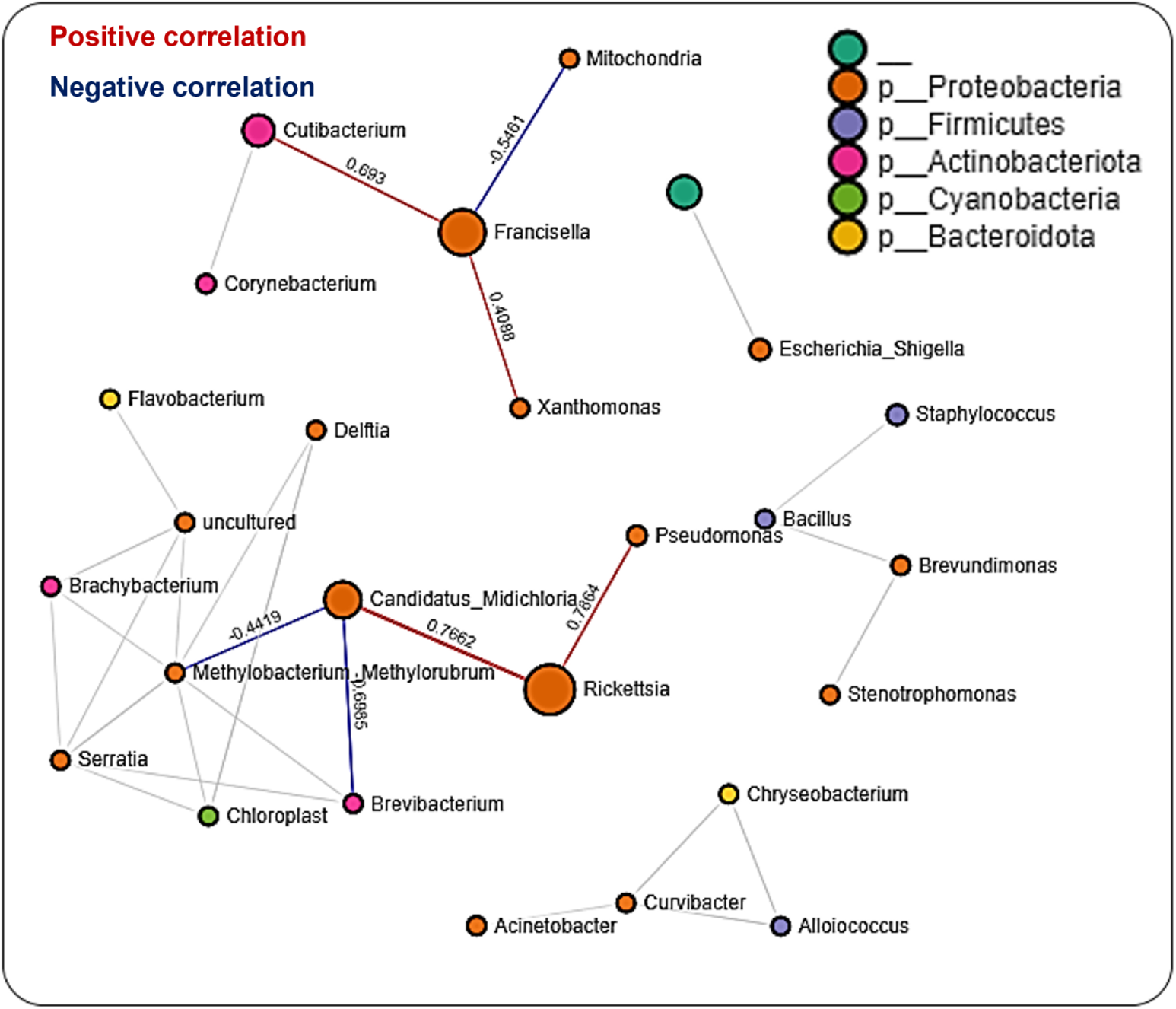
Correlation network analysis in the midgut of *Am. maculatum* ticks. Correlation network generated using the SparCC algorithm. Correlation network with nodes representing taxa at the family level and edges representing correlations between taxa pairs. Node size correlates with the number of interactions in which a taxon is involved, and the line color represents either positive (red) or negative (blue) correlations. The color-coded legend shows the bacterial family to which each taxon belongs.

### Microbial interaction in the midgut

We observed many fewer (56) significant partial correlations in the midgut tissues between 29 bacteria genera, all belonging to the phyla Proteobacteria (15 genera), Firmicutes (3 genera), Actinobacteriota (4 genera), Cyanobacteria (1 genus), and Bacteroidota (2 genera) as well as an unknown phylum (1 genus), 16 (29%) of which were negative correlations and the remaining 40 (71%) positive correlations (Additional File, Table 7). The network map of bacterial interactions identified five distinct networks, with *Rickettsia* and *Candidatus*_*Midichloria* observed to be part of the same network with positive correlations. *Francisella* was found to be in a different network (Fig. 6). A striking observation was the ubiquitous presence of bacteria in the phylum Proteobacteria across all five identified networks (Fig. 6). The LDA score from LEfSe identified the enrichment of *Rickettsia, Escherichia_Shigella,* and *Cutibacterium* in both uninfected and *R. parkeri-*infected midgut tissues. Compared with the *R. parkeri-*infected midgut, only *Francisella* was significantly enriched in the uninfected midgut, while *Bacillus* was enriched in the *R. parkeri-*infected midgut (Additional File, Fig. 6A). Assessing the effect of blood meal on bacterial enrichment identified a shared enrichment of *Rickettsia, Escherichia_Shigella,* and *Cutibacterium* in both unfed and partially fed midgut. We observed *Pseudomonas* and *Brevibacterium* in the unfed midgut, while *Francisella* and *Staphylococcus* were enriched in partially blood-fed midgut tissues (Additional File, Fig. 6B). Microbiota– microbiota interactions across developmental stages were driven mainly by three of the most abundant bacteria: *Francisella, Rickettsia,* and *Candidatus_Midichloria*. Other interactions revealed that certain taxonomic groups were inversely correlated, as seen between the phyla Firmicutes and Proteobacteria. Many specific interactions were observed in the tissues, suggesting tissue-dependent influences on how microbes interact in different tissue niches.

### Functional characterization of bacterial communities

Functional prediction of microbiota genes compared with COG functional categories identified genes involved in ribosomal structure and biogenesis, nucleotide transport, and metabolism as the most abundant across developmental stages (Additional file, Table 8). Several genes involved in metabolisms, such as lipid transport, coenzyme transport, amino acid transport, and carbohydrate transport, were also identified (Additional files, Fig. 10). Prediction of metabolic genes and pathways using the KEGG Orthology (KO) categories assigned 19% of the metabolic functions to genes in carbohydrate metabolism pathways and 18% of the functions were encoded by genes in amino acid metabolism pathways. By comparison, genes encoding cofactors in the vitamin metabolism pathways and energy metabolism pathways represented 15% and 13% of the metabolic pathways, respectively (Additional file, Fig. 10).

Differences in functional community profiles within the metabolic pathways were further examined between the different feeding stages and infection status across dissected tissues based on the KEEG metabolism categories. The partially fed salivary gland and midgut had a higher proportion of sequences assigned to energy metabolism and carbohydrate metabolism than the unfed salivary gland and midgut (Fig. 7A, B). The presence of *R. parkeri* led to an increase in the proportion of glycan biosynthesis and metabolism genes, carbohydrate metabolism genes, and genes involved in the metabolism of cofactors and vitamins in the salivary glands. A slight increase was also seen in genes involved in the metabolism and biodegradation of xenobiotics in the *R. parkeri-*infected salivary gland (Fig. 8A). A slight increase in the proportion of energy metabolism genes was associated with *R. parkeri* in infected midgut compared with the uninfected midgut (Fig. 8B). Interestingly, genes involved in amino acid metabolism were seen to be reduced in *R. parkeri-*infected salivary glands and midgut compared with uninfected tissues (Fig. 8A, B). No differences between metabolic predictions were detected between *R. parkeri-*infected and uninfected ovarian tissues (Additional file, Fig. 11).

**Figure 7:**
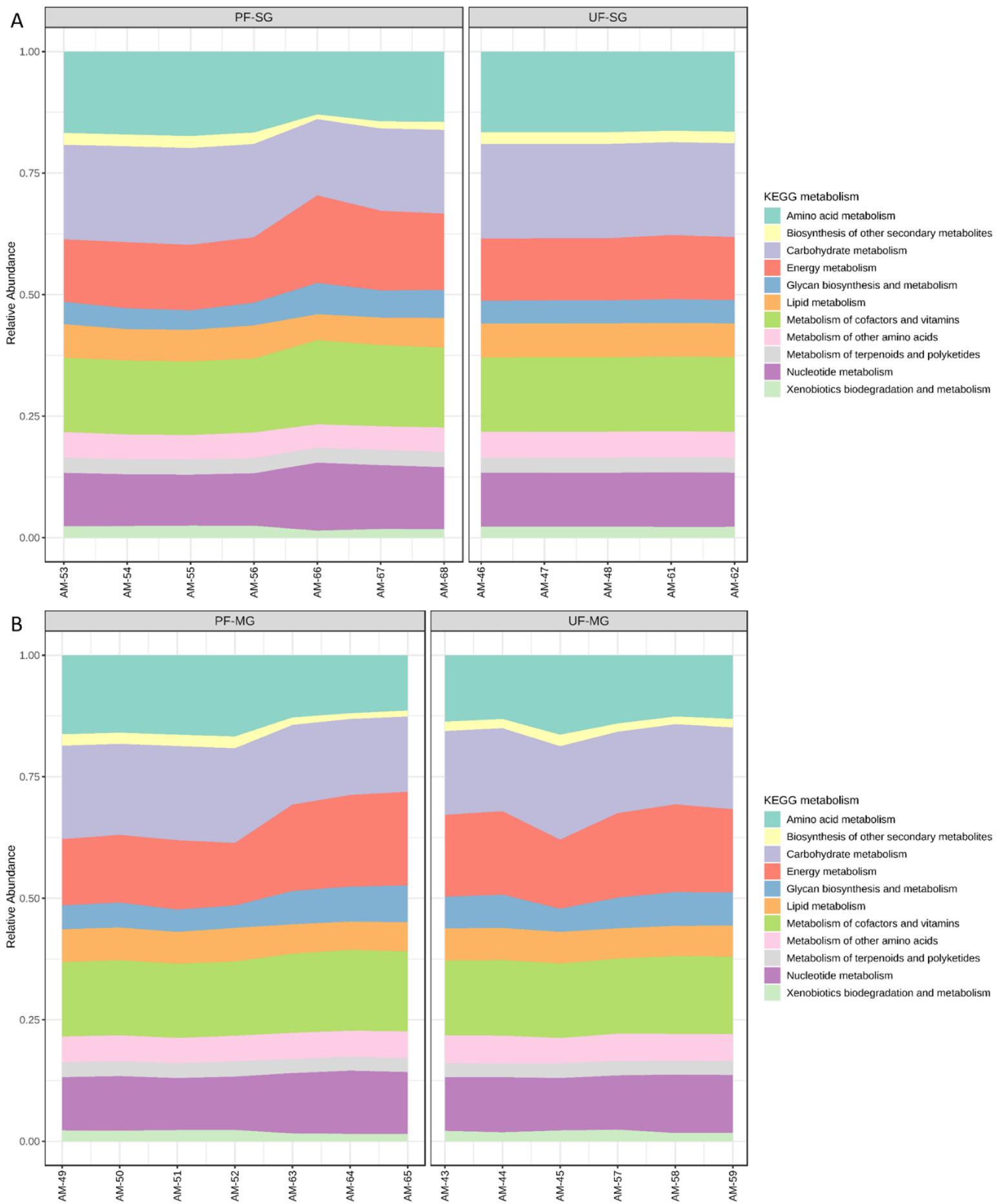
Histogram indicating the functional differences between the fed and partially fed *Am. maculatum* microbiota in A) salivary gland and B) midgut tissues. KEGG metabolic categories were obtained from 16S rRNA gene sequences using the PICRUSt2 pipeline.

**Figure 8:**
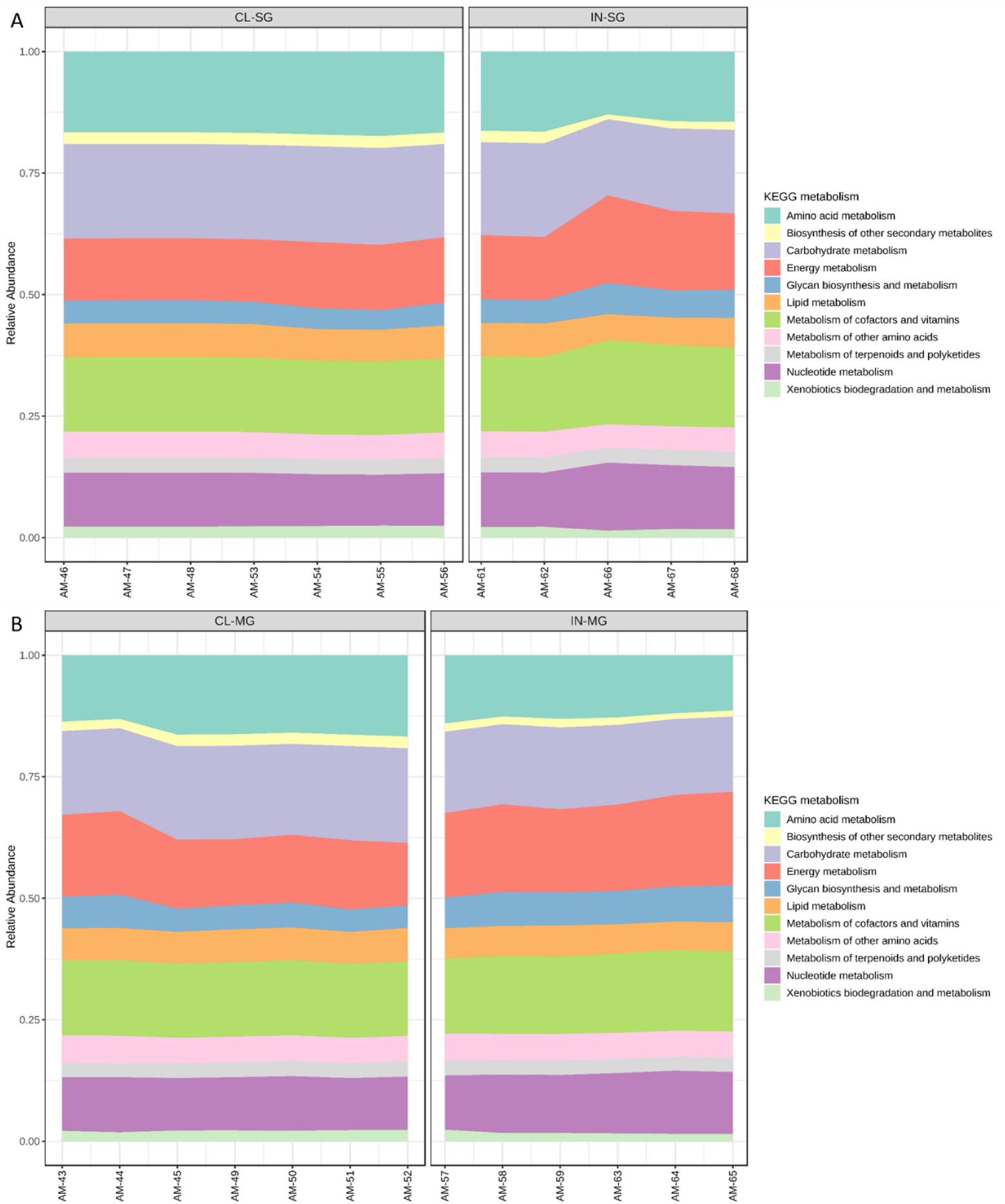
Histogram indicating the functional differences between the uninfected and *R. parkeri*-infected *Am. maculatum* microbiota in A) salivary gland and B) midgut tissues. KEGG metabolic categories were obtained from 16S rRNA gene sequences using the PICRUSt2 pipeline.

## Discussion

This study is the first to characterize microbiome changes throughout the ontogeny of the Gulf Coast tick, a competent vector of the spotted fever group rickettsiae, *R. parkeri*. Using a high-throughput metagenomic approach that allowed us to barcode DNA extracted from individual ticks and process them in a single run, the present study set out with the aim of assessing the core microbial assemblages present in *Am. maculatum* ticks and determine the changes under certain physiological conditions vis-à-vis development, hematophagy, and pathogen infection. The results show that changes to the core microbiome occur in response to a blood meal and pathogen association but not as a function of the developmental stage. Similarly, as expected, microbe-microbe interactions depended on the specific tick tissues in which those interactions took place.

### The *Amblyomma maculatum* microbiome remains stable across ontogeny

The typical life cycle of *Am. maculatum* ticks encompass critical physiological and environmental events conferred by intrinsic and extrinsic factors, such that the persistence of the microbiome and every similar micro-community that utilizes the tick as a microenvironment across these dynamic events is continuously tested. Two very well-studied factors that exert significant pressure on the tick microbiome are blood meal and pathogen replication (Abraham et al., 2017; Swei and Kwan. 2017; Adegoke et al., 2020; Wei et al., 2021), albeit with varying outcomes reported. In the current study, a shift favoring the establishment of a single bacterial genus along developmental lines (eggs to adult male and female) is exhibited (Fig. 1A–C). Even so, the uptake and breakdown of nutritious blood meals and the intricacies of tick–pathogen interactions involve several detailed mechanistic processes that have been shown to significantly impact tick physiology (de la Fuente et al., 2016; Hoxmeier et al., 2017; Budachetri et al., 2018), the current study shows that members of the *Am. maculatum* microbial communities exhibit reduced plasticity thus maintaining a stable microbial core. This observation raises the question of whether the tick microbiome is self-regulated in ticks and tick tissues

Notably, while the core microbiome persists across developmental stages, certain bacterial genera were found to be associated with both blood meal and pathogen infection, as seen with *Cutibacterium* and *Candidatus*_*Midichloria*, and whether the introduction of these bacteria in fewer numbers significantly impacts overall tick biology is a question beyond the scope of this study (Budachetri et al., 2018; Adegoke et al., 2020, Abraham et al., 2017). To further understand how changes across tick developmental stages shape microbial assemblages, it is imperative to assess microbial assemblages and community measures of diversity, such as richness and evenness (Brown et al., 2020). Community diversity is important because the smallest differences at the taxon level induced by blood meal or pathogen interaction could interfere with community structure, which would otherwise not be reflected in the abundance profile. We observed a continuous decrease in community diversity towards later developmental stages (Figure 1D–I). For both diversity metrics employed, a reduction in community richness likely indicated a corresponding decline in all or certain specific bacterial taxa. At the same time, a decrease in evenness would imply that certain bacterial taxa are allowed to increase in abundance at the expense of other bacteria. This finding is consistent with observations made in the kissing bugs, where there is a decrease in diversity associated with ontogenic development (Mann et al., 2020), which is in disagreement with Oliveira et al. (2018), who reported an increase in diversity with ontogenic development. A possible explanation for this discrepancy may be attributed to internal and external changes that occur during metamorphosis. Shedding of the midgut lining and the external cuticles are two of the most important phenomena during development. These may come with significant loss of resident and non-obligate bacteria whose absence poses no detrimental effect on tick biology (Philipp et al., 2013). There are, however, other possible explanations for the reduction in diversity seen in this study.

One such explanation is that bacteria that are not vertically maintained from the female to her eggs are replaced by vertically maintained bacteria. The preservation of vertically maintained species would explain the loss of *Cutibacterium* in adult male and female ticks. By extension, there would be a reduction in overall diversity. Previous findings have highlighted the importance of initial exposure to bacteria in eliciting a robust immune response upon subsequent exposure to the same bacteria or bacteria from a similar group, a phenomenon referred to as immune priming (Trappeniers et al., 2019; Powers et al., 2020). This could explain the fact that immature developmental stages of ticks are easily infected with bacteria from the environment because of their naïve immune status. These bacteria will not be carried over to the adult stage due to a more developed immune status. The persistence of *Francisella, Rickettsia,* and *Candidatus*_*Midichloria* to the adult male and female further emphasizes their position as candidates for vertical maintenance, as previously reported for bacterial species from the same genus (Baldridge et al., 2009; Wright et al., 2015; Azagi et al., 2017; Mukhacheva and Kovalev, 2017; Budachetri et al., 2018).

### Microbial assemblages in tissues are driven by pathogen and host blood

To gain in-depth insight into how different arthropod tissues contribute to tick biology and, most importantly, their role in vector competence, it is imperative to know how resident microbial communities differ from one tissue to the other. Several factors have been identified that significantly affect how the microbiome is shaped in different arthropod tissues, albeit with varying outcomes and findings. In this study, we asked whether differences in microbial composition occur within tissues and whether changes influence these differences in the state of blood-feeding and infection status. Our results clearly show a difference in the abundance and composition of the bacteria identified across tick tissues. A fascinating conclusion from our findings is the higher bacterial proportion in saliva, which does not correspond to the bacterial balance in the salivary glands. Tick saliva contains simply a snapshot of the microbes secreted by the tick and is possibly not a good representation of all microbes delivered during prolonged feeding. These differences can be explained in part by the fact that some of the microbial populations identified in the saliva, such as *Bacillus* and *Staphylococcus,* are known environmental bacteria, have also been reported to be found on the skin and fur of the animal hosts (Dastgheyb and Otto, 2015; Onyango and Alreshidi, 2018; Ravine, 2019), and may have been acquired during the feeding process. Some of these bacteria could also have been mechanically transferred during saliva collection from the ticks. The presence of the microbiome within tick saliva, while preliminary, could implicate the tick microbiome in modulating the host immune response to tick bite at the host–tick interface. The observation of a higher diversity seen in midgut tissues compared with salivary gland, ovary, and saliva contrasted with the findings of Zolnik et al. (2016), who reported a higher diversity in salivary gland compared with midgut tissue. The tick midgut and associated peritrophic membrane play an important role in blood breakdown and its components by several catalytic proteins. The production of toxic reactive oxygen species (ROS) and reactive nitrogen species (RNS), which follow blood digestion, is also dampened by these midgut proteins, thus enabling midgut-resident microbial replication. The heme component of the blood meal derived by catalytic proteins also counters the potential tissue damage associated with a heme-induced oxidative burst. Another plausible explanation for the higher microbial diversity observed in the midgut reported from this study is that the midgut is the only tissue that directly encounters host blood, and it provides the first line of contact with any bacterium coming in with the blood meal (Houk, 1977; Tarnowski and Coons, 1989; Gaio et al., 2011; Lejal et al., 2019). However, prolonged blood feeding and, by extension, the initiation of blood meal digestion might provide an enabling environment for bacterial replication and growth, hence impacting the dynamic and increasing overall diversity of the midgut microbiome as seen from this study. Due to the importance of the ovarian tissues in developmental success, it is expected that several physiological barriers will be put in place to confer protection and reduce the tick’s interaction with circulating microbes to the barest minimum. The reduced microbial diversification observed in the ovary is consistent with this, as seen in Fig. 2D. Several studies have outlined the contribution of invertebrate hemocytes to immune functions while also identifying distinct populations of hemocytes as either circulating or sessile (Kadota et al., 2003; King et al., 2013; Sigle and Hillyer, 2018). The sessile population of hemocytes is usually found attached to cavity walls, such as the abdomen, thorax, and internal organs. While extensive studies have described how arthropod hemocytes play a significant role in immunomodulation (Pereira et al., 2001; Hillyer, 2009; Liu et al., 2011; Hillyer and Strand, 2014; Fiorotti et al., 2019; Kwon et al., 2019), few studies have defined the functional role of sessile populations in the biology of the organism (Sigle and Hillyer, 2016). Hence, it could be hypothesized that circulating or sessile hemocytes would form immunological barriers as part of an extensive physiological barrier to protect a vital organ, such as the ovary, as seen with several human and mammalian blood–tissue barriers (Fröhlich, 2002)

It has already been established that blood uptake, which usually lasts several days in ticks, can introduce a plethora of pathogenic and non-pathogenic microbes into ticks (Bhatia et al., 2018; Zolnik et al., 2018; Landesman et al., 2019). While ticks acquire host blood for nutritional requirements, several host factors present in the blood meal could also potentially contribute to compositional changes that are likely to occur to the microbiome throughout the extended feeding process. For instance, the tick midgut responds to the presence of the blood meal via intricate protein expression patterns (Rachinsky et al., 2008; Schwarz et al., 2014; Perner et al., 2016; Oleaga et al., 2017; Liu et al., 2018) while also undergoing certain morphological and structural adjustments (Tarnowski and Coons, 1989; Frants et al., 2010), all of which could have an impact on the microbial environment in the midgut, as previously reported (Zhang et al., 2014; Zolnik et al., 2018; Landesman et al., 2019). Results from our study show that microbial diversity is impacted following uptake of the blood meal in both salivary gland and midgut tissues (Fig. 2E). We also observed a combinatorial effect of blood meal and *R. parkeri* infection in reducing the abundance of *Francisella* while increasing *Rickettsia* abundance in the partially fed and infected salivary gland and, vice-versa, in the partially fed and infected midgut. This suggests that there is the trafficking of *Rickettsia* from the midgut into the salivary gland during feeding. While studies that assessed the impact of blood meal on the tick microbiome are limited to developmental stages, it is interesting to note that our studies corroborate a great number of these studies, albeit in life stages. Zolnik and colleagues (2016) observed a significant reduction in alpha-diversity throughout the feeding period of nymphal ticks compared with the unfed nymphs of black-legged ticks, and Zhang et al. (2014) also reported an altered microbial composition following a prolonged blood meal in *Ixodes persulcatus* ticks. A similar observation was reported in the malaria mosquito, in which Muturi et al. (2019) found a significant decrease in microbial diversity in blood-fed *Aedes aegypti* compared with sugar-fed *Ae. aegypti* mosquitoes. While comparing the effect of different host blood meal sources was beyond the scope of this study, our findings strongly implicate host blood meal as a significant factor in driving tick microbiome diversity, which may pose a significant effect on tick biology, as previously shown by Swei and Kwan, 2016.

### Microbe–microbe interactions are driven by the specific host tissue and not the developmental stage

#### Microbial interactions across developmental stages

The biological success of the tick as a hematophagous arthropod and a vector of disease pathogens hinges partly on the specific interactions between different members of the resident microbiota, which encompasses the entirety of all the microorganisms present in an environment. Identifying unique interactions between these microbial community members is a critical step in developing effective control measures against tick-transmitted pathogens.

Our network analysis on the overall dataset across all developmental stages identified similar percentages of positive and negative correlations, suggesting a much more balanced distribution of both antagonistic (Budachetri et al., 2018) and synergistic (Eiler et al., 2012; Chow et al., 2014) interactions between members of the microbial communities. This observation contrasts with a recent study by Lejal et al. (2021), who reported more than 97% partial positive correlations across all the datasets in *Ixodes ricinus* ticks and suggested that most of the tick microbial community favors a mutualistic interaction. While the findings of our study contrast with this observation, it should be stated that microbial correlations of an individual organism may not fully represent interactions in different tissue niches, which are often a microenvironment for unique bacteria communities (Pollet et al., 2020). The genera *Rickettsia*, *Francisella,* and *Candidatus*_*Midichloria* were all part of the same network, with positive correlations between all three bacterial genera, and these were previously identified in *Am. maculatum* ticks (Budachetri et al., 2014; Budachetri et al., 2018) and *Ix. ricinus* ticks (Lejal et al., 2021). At the same time, *Francisella* and *Candidatus*_*Midichloria* are major endosymbionts found in the *Am. maculatum* microbiome and Budachetri et al., (2018) reported an increase in *Candidatus*_*Midichloria* load following *R. parkeri* infection, proposing a positive correlation between these bacteria and a corresponding reduction in *Francisella* load following *R. parkeri* infection, implying a negative correlation.

Interestingly, recent network analysis on the *Ix. ricinus* microbiota also shows a positive correlation between *Rickettsia* and *Candidatus*_*Midichloria* OTUs, indicating a facilitative interaction between these bacteria (Lejal et al., 2021). Our data suggest a positive interaction between the pathogenic Rickettsia and the obligate endosymbiont *Candidatus*_*Midichloria*. The current study’s findings are consistent with those of Budachetri et al. (2018), who reported a synergistic interaction between *R. parkeri* and *Candidatus*_*Midichloria mitochondrii* and an antagonistic interaction between *R. parkeri* and a *Francisella*-like endosymbiont. While identification at the species level exceeded the resolution of our sequencing analysis, we have strong reason to believe that the OTU belonging to the *Coxiella* genus is the *Coxiella*-like endosymbiont (CLE), because the ticks used in this study were strictly maintained under laboratory conditions. CLE is ubiquitously present in most hard ticks, with reports suggesting an obligate association with the tick host, in which they provide essential vitamin supplements for tick physiological success (Duron et al., 2015; Smith et al., 2015; Guizzo et al., 2017; Ben-Yosef et al., 2020).

A recent study reported the replacement of bacteria in the *Coxiellaceae* family with those in the *Francisellaceae* family in both lab-maintained and field-*collected Am. americanum* ticks (Kumar et al., 2021), providing further evidence of competition between these two tick endosymbionts. The strict adherence to a single nutritional mutualist by tick species is an area of tick–microbial interaction that requires further exploration, and future studies should aim at characterizing both the immunoproteome and metabolites of gnotobiotic ticks with known obligate endosymbionts. Other key interactions observed between the OTUs belong to the *Bacillus*, *Rickettsia,* and *Francisella* genera. The genus *Bacillus* is an environmental resident microbe that ticks acquire while questing in the environment and during the attachment process to the host (Ross et al., 2019). We observed negative correlations between *Bacillus* and *Rickettsia* and between *Bacillus* and *Francisella,* suggesting a competitive interaction between these bacterial OTUs. The findings of the current study are consistent with those of another study (Lejal et al., 2021) that described a positive correlation between *Bacillus* and *Rickettsia* but was not compatible with the presence of a high abundance of *Bacillus* in *Theileria-*positive *Rhipicephalus microplus* ticks, as previously described (Adegoke et al., 2020). The factors that favor the establishment and long-term association of environmentally acquired bacteria by ticks still need to be experimentally identified. Whether these bacteria express effector molecules that suppress pathogenic microbes, such bacteria may be utilized as effective vector-control tools in the future to develop paratransgenic ticks.

### Microbial interactions are driven by the specific tissue niche

Network analysis on datasets representing all tissues was initially carried out, thereby identifying 67% positive correlations between different bacteria in the networks. We identified more positive correlations across tissue levels than developmental stages, suggesting a much finer association scale at smaller niches (Pollet et al., 2020). Aside from the higher proportion of positive correlations, most of the microbial networks identified when applied to the combined tissue dataset were like those presented for developmental stages. For instance, OTUs from *Francisella, Rickettsia, and Candidatus_Midichloria* were all present in the combined tissue networks, including the salivary gland and midgut. Interestingly, while the salivary gland dataset had a relatively higher proportion of networks, it contained more positive correlations (71%). Only OTUs belonging to the genera *Francisella, Rickettsia,* and *Candidatus*_*Midichloria* were involved in most interactions. An unexpected outcome from the salivary gland network was the observation of a negative correlation between *Rickettsia* and *Candidatus*_*Midichloria* OTUs, while there was no direct correlation between *Francisella* and either of *Rickettsia* or *Candidatus*_*Midichloria*. We proposed that *Candidatus*_*Midichloria* in the salivary gland was not transmitted to the host, due to its restricted location in the mitochondria, hence its absence from the saliva. This phenomenon was also observed in the midgut, albeit with fewer and more distinctly separated microbial networks. A handful of studies deciphered specific correlations between microbial members residing in arthropod vectors (Lejal et al., 2021; Hedge et al., 2018). While our findings here are preliminary at best, they emphasize that tissue-specific microbial interactions do exist, and this should be further investigated, as they may lead to new insights into pathogen replication and establishment across tick tissues. Future work should be targeted towards experimental validation of the predicted correlations to understand how these interactions are shaped.

The metabolic predictions based on the microbiota showed no differences in metabolic genes across developmental stages; however, we identified a diverse category of genes involved in metabolic pathways, most of which have been previously reported in ticks and other blood-feeding arthropods (Obregon et al., 2019; Bahrndorff et al., 2018; Jing et al., 2020; Couper et al., 2019). The presence of energy metabolism genes and pathways in higher abundance in the partially fed than the unfed salivary gland and midgut tissues did not completely align with the microbial changes induced by blood meal, especially with the midgut. We saw a significant increase in the microbial profiles in the partially fed salivary gland compared with the unfed salivary gland (Fig. 2B). We hypothesize that since the increased proportion of bacterial genes was involved in energy metabolism, carbohydrate metabolism could be a physiological pressure on the salivary gland as a secretory organ involved in blood meal at the host–feeding interface. The slight increase in the number of genes involved in xenobiotic degradation and metabolism in the partially fed salivary gland could be due to the constant need to counter host immune responses to achieve hematophagy.

Interestingly, metabolic pathways involved in energy metabolism and the metabolism of vitamins and cofactors were upregulated in *R. parkeri-*infected salivary glands and midgut tissues (Fig. 8A, B). These changes were also similar to those associated with uptake of blood meal. While we could not tease out whether these changes were solely associated with either blood meal or *R. parkeri* infection, we saw a slight increase in the abundance of genes involved in nucleotide metabolism pathways in the *R. parkeri-*infected salivary gland. The importance of nucleotide metabolism pathways in bacteria is highlighted by their role in bacterial physiological processes and the synthesis of new nucleic acids necessary for replication (DNA) and transcription (RNA) (Lopatkin and Yang, 2021). Our current results show that blood meal increases the relative abundances of *Francisella* and *Candidatus_Midichloria* endosymbionts. Likewise, the abundance of *Rickettsia* also increases with *R. parkeri* infection, suggesting an overall increase in the replication of these bacterial groups, thus leading to an overall increase in nucleic acid replication and transcription. While our functional analysis is predictive at best, with approximately 85% similarity with shotgun metagenomics, it did provide comprehensive insight into the different functional pathways present in the tick microbiota.

## Conclusion

The microbiome of *Am. maculatum* comprises ten bacterial genera. Three genera, including *Rickettsia, Francisella*, and *Candidatus_Midichloria,* predominate in the core microbiome. Blood-feeding behavior aided in the proliferation of these three bacteria, whereas overall diversity and evenness were reduced due to elevated levels of ROS associated with blood meal acquisition and digestion, both in whole ticks and in particular organs (gut/salivary glands/ovaries). *R. parkeri* infection co-proliferates with *Candidatus_Midichloria,* while an antagonistic interaction with the *Francisella* genus was observed in the tick life cycle. *R. parkeri* infection further increased the diversity or evenness of the core microbiome. A significant increase in activity of the genes related to metabolic pathways was observed in ticks after a blood meal and pathogen infection. With these results in mind, further study should focus on the interplay between these three bacterial species, including the pathogen, while studying tick–pathogen interactions. The interplay of symbiotic bacteria (*Candidatus_Midichloria/Francisella*) with pathogenic *R. parkeri* should yield important findings in future studies.

## Acknowledgments

The authors thank Surendra Raj Sharma, Latoyia Downs, and Faizan Tahir for thought-provoking and enriching discussions, particularly about endosymbionts, reproductive fitness, and priming of the tick immune system. Finally, we are grateful to Dr. Michael Garrett, UMMC genomics core facility.

## Declarations

### Funding

This research was principally supported by the USDA NIFA award #2017-67017-26171, the Pakistan–US Science and Technology Cooperation award (US Department of State); the NIH NIAID award #R15AI099910; the NIH NIGMS award #R15GM123431; and the Mississippi INBRE (an institutional award (IDeA) from the National Institute of General Medical Sciences of the National Institutes of Health award #P20GM103476). The funders played no role in the study design, data collection, analysis, publication decision, or manuscript preparation.

### Author Information

**School of Biological, Environmental, and Earth Sciences, University of Southern Mississippi, Hattiesburg, MS 39406, USA**

Abdulsalam Adegoke, Deepak Kumar, Khemraj Budachetri, Shahid Karim

**Center for Molecular and Cellular Biosciences, University of Southern Mississippi, Hattiesburg, MS 39406, USA**

Shahid Karim

## Abbreviations

Am. maculatum: Amblyomma maculatum

PCA: Principal component analysis OTU: Operational taxonomic unit TBD: Tick-borne disease

TBP: Tick-borne pathogens

FLE: Francisella-like endosymbiont

CMM: Candidatus Midichloria mitochondrii

QIIME2: Quantitative Insights into microbial ecology 2 SparCC: sparse correlations for compositional data LEfSe: linear discriminant analysis effect size

LDA: linear discrimination analysis

## **Et**hical Approval and Consent to participate

Not applicable.

## Consent for publication

Not applicable

## Availability of data and materials

Raw sequences for this study were submitted to the NCBI read under SRA database and an accession # obtained.

## Competing interests

The authors declare that they have no competing interests.

## Author contributions

Conceptualization: Shahid Karim

Data curation: Abdulsalam Adegoke, Deepak Kumar, Shahid Karim Formal analysis: Abdulsalam Adegoke, Deepak Kumar, Shahid Karim Funding acquisition: Shahid Karim

Investigation: Abdulsalam Adegoke, Shahid Karim

Methodology: Abdulsalam Adegoke, Deepak Kumar, Khemraj Budachetri Project administration: Shahid Karim

Resources: Shahid Karim Supervision; Shahid Karim

Validation: Abdulsalam Adegoke, Shahid Karim Visualization: Abdulsalam Adegoke

Writing, original draft: Abdulsalam Adegoke, Shahid Karim

Writing, review & editing: Abdulsalam Adegoke, Deepak Kumar, Khemraj Budachetri, Shahid Karim

## ADDITIONAL FILES

**Figure S1:**
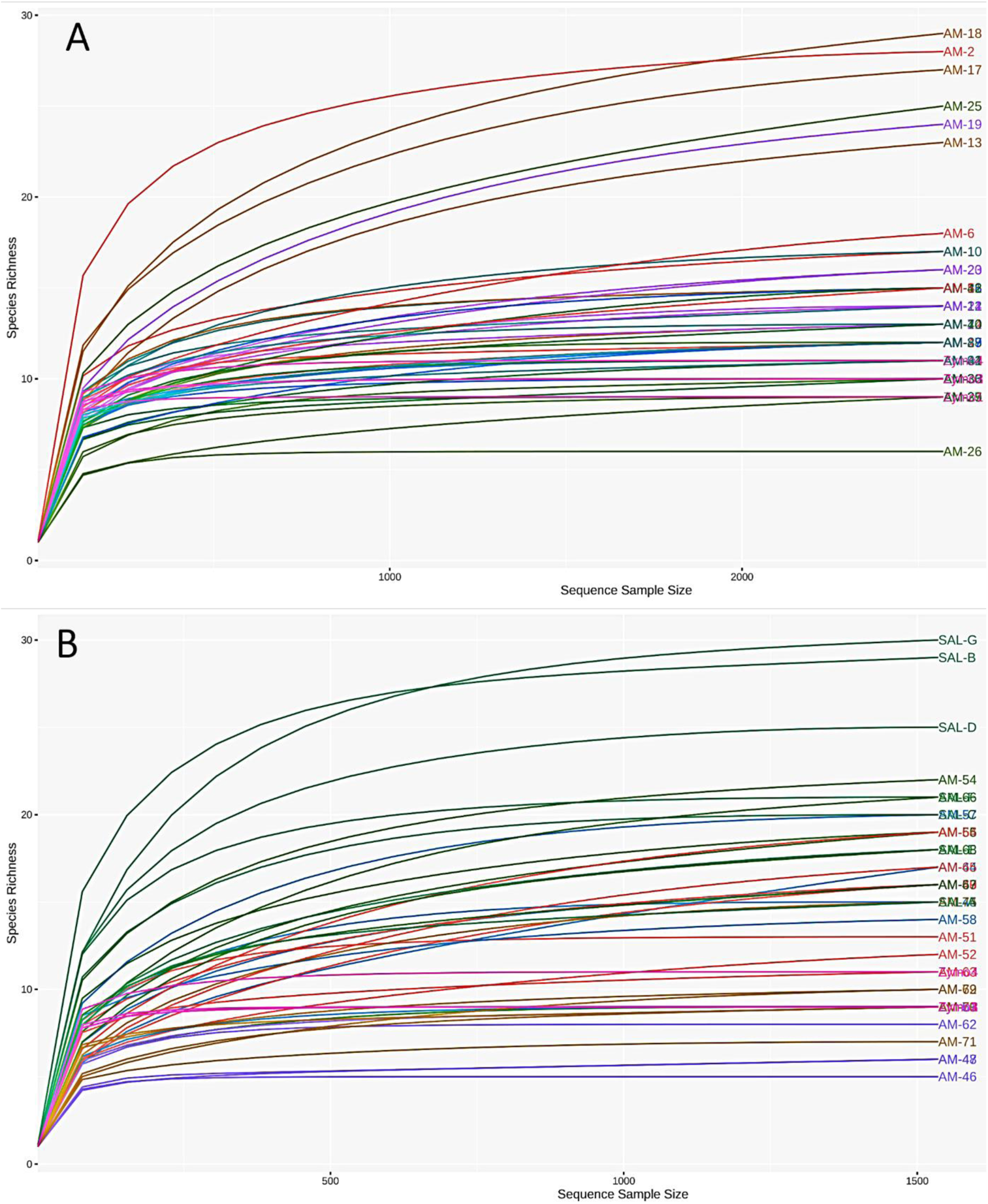
Rarefaction analysis of raw reads from A) developmental stages and B) isolated tissues of *Am. maculatum* ticks. Each curve representing individual replicates was rarified to a sequence depth of approximately 3000 sequences for all developmental stages (unfed and partially fed) and 1000 sequences for isolated tissues.

**Figure S2:**
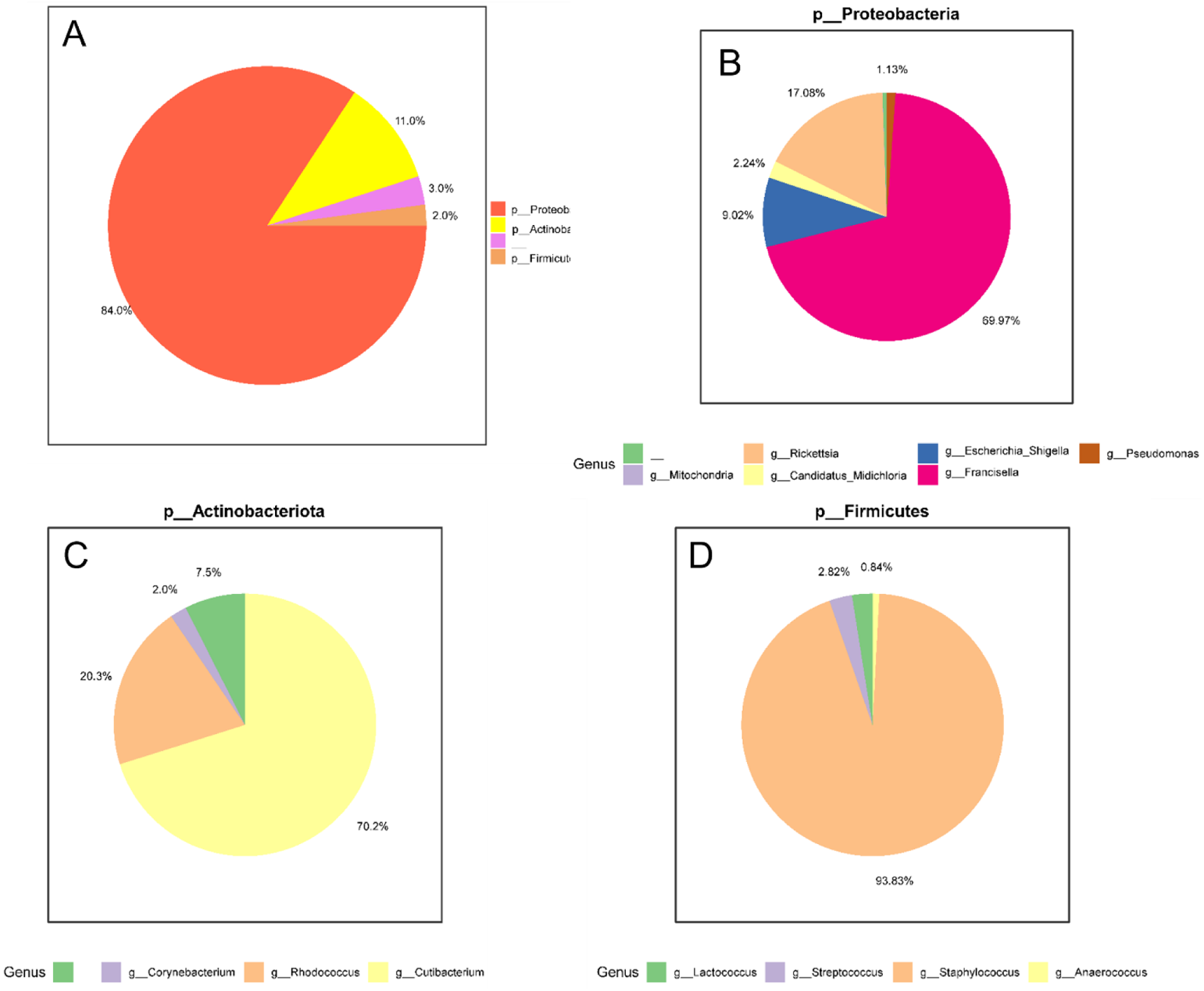
Pie chart summary of microbial abundances in the developmental stages of *Am. maculatum*. A) Overall abundance summary at the phylum taxonomic level and breakdown of abundant phyla showing B) Proteobacteria, C) Actinobacteriota, and D) Firmicutes. A representative genus from each phylum is indicated in the legends.

**Figure S3:**
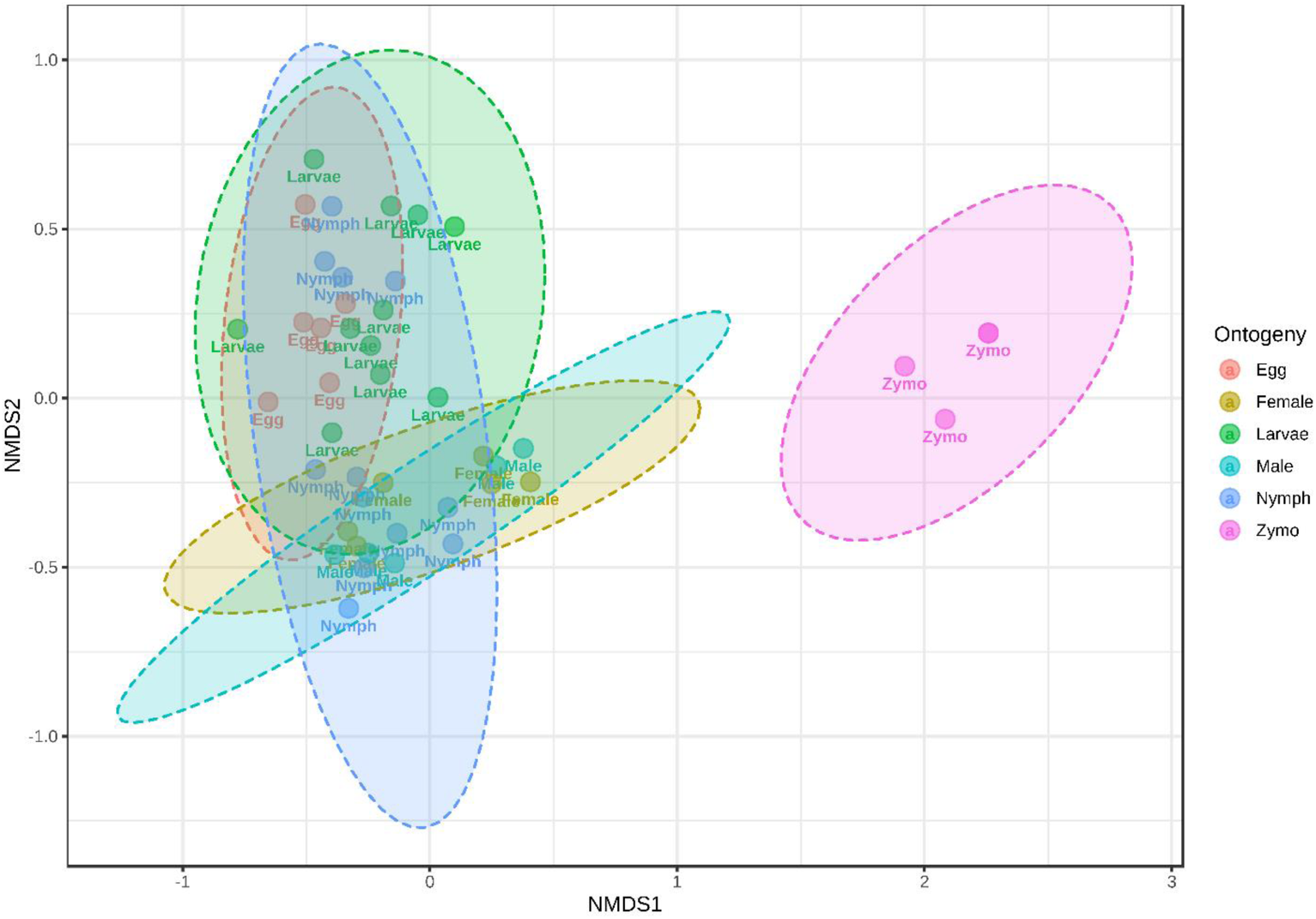
Non-metric multidimensional scaling (NMDA) analysis of β-diversity measures across *Am. maculatum* developmental stages. Ellipses around unique clusters indicate the degree of distance based on the Bray–Curtis distance matrix.

**Figure S4:**
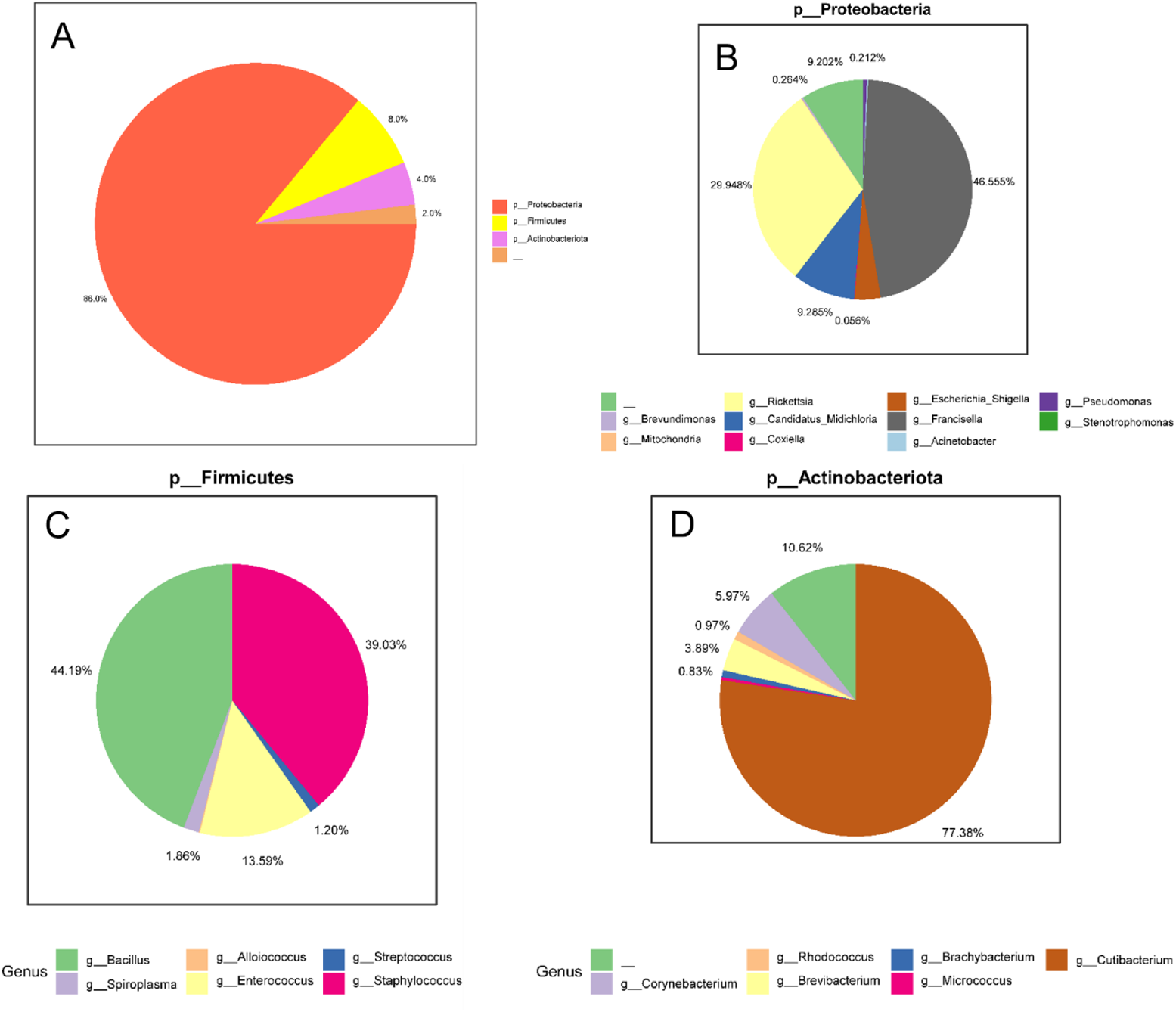
Pie chart summary of microbial abundances in the dissected tissues of *Am. maculatum*. A) Overall abundance summary at the phylum taxonomic level and breakdown of abundant phyla showing B) Proteobacteria, C) Firmicutes, and D) Actinobacteriota. A representative genus from each phylum is indicated in the legends.

**Figure S5:**
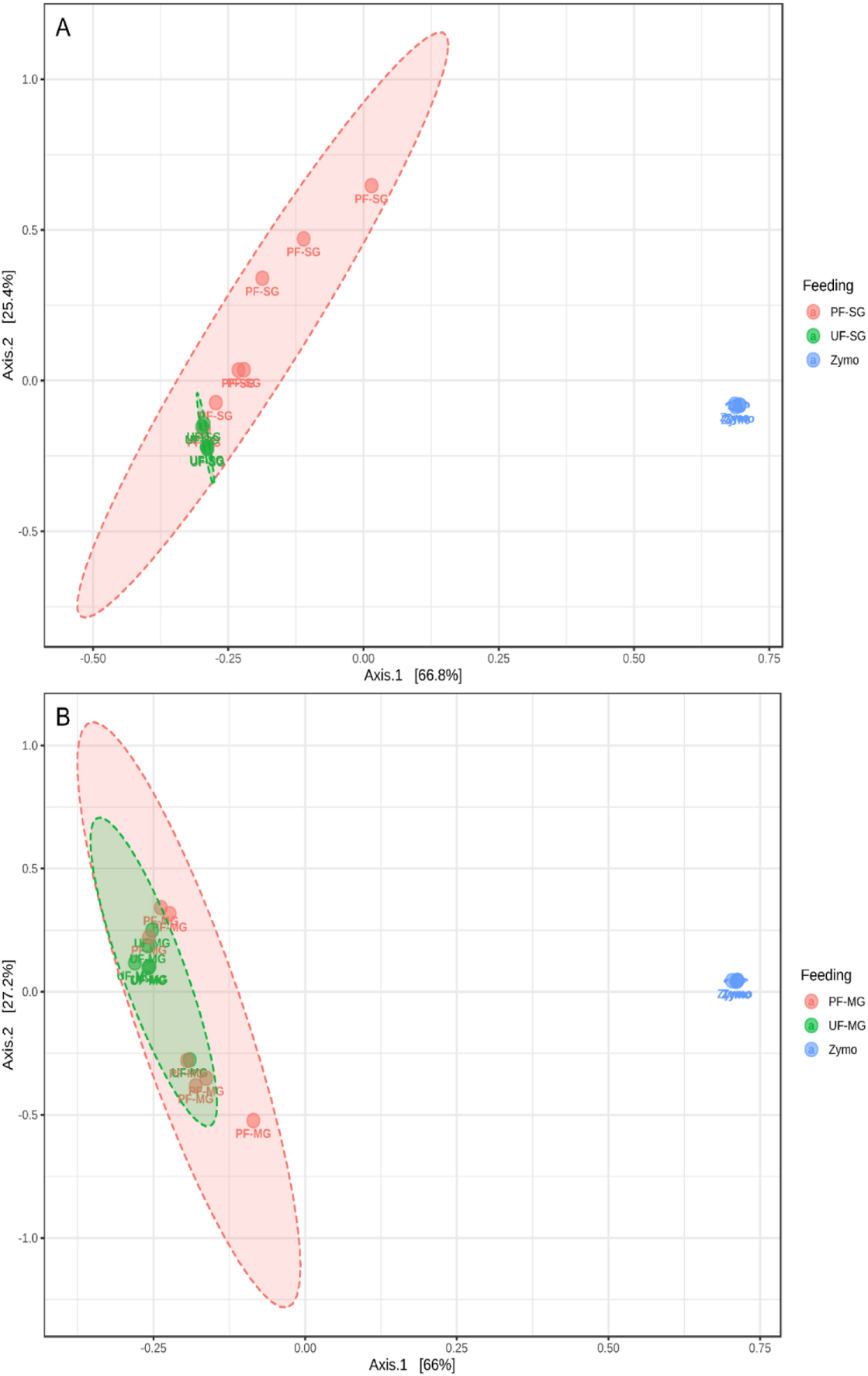
Principal coordinate analysis of β-diversity measures of tissues dissected from unfed and partially fed *Am. maculatum* ticks. Bray–Curtis distance matrix of A) salivary gland and B) midgut tissues. Ellipses around unique clusters indicate the degree of distance.

**Figure S6:**
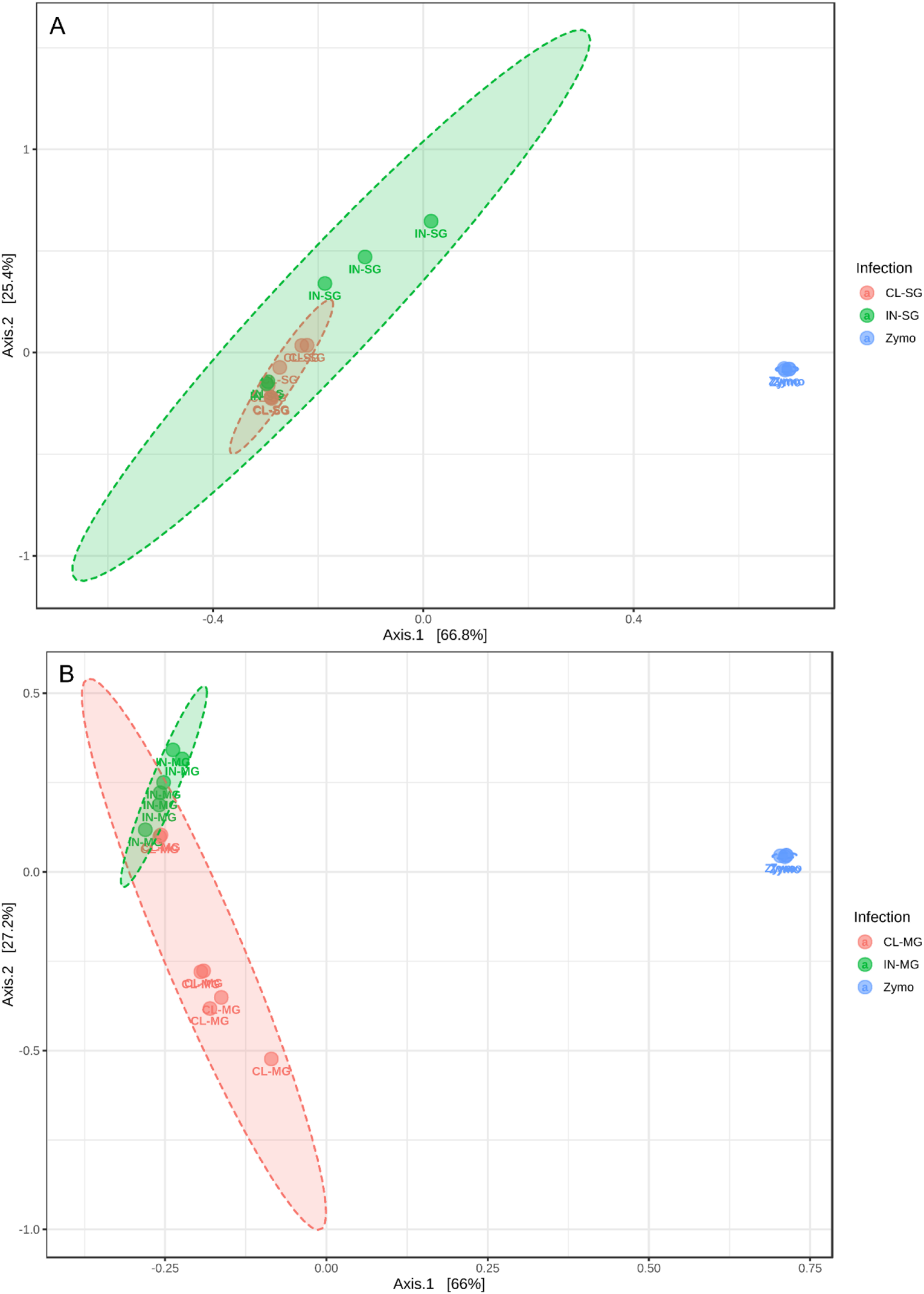
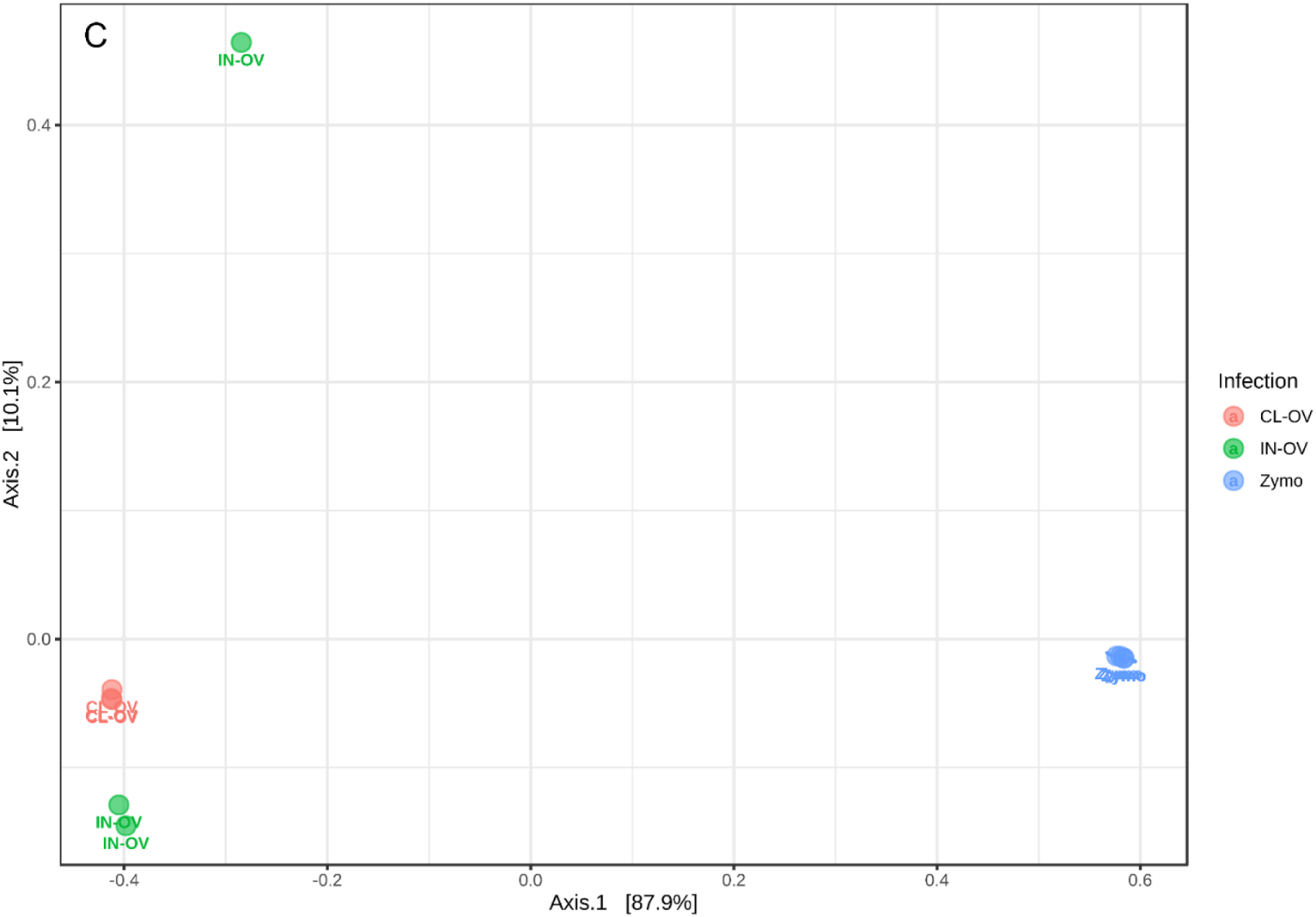
Principal coordinate analysis of β-diversity measures of tissues dissected from unfed and partially fed *Am. maculatum* ticks. Bray–Curtis distance matrix of A) salivary gland, B) midgut tissues, and C) ovarian tissues. Ellipses around unique clusters indicate the degree of distance.

**Figure S7:**
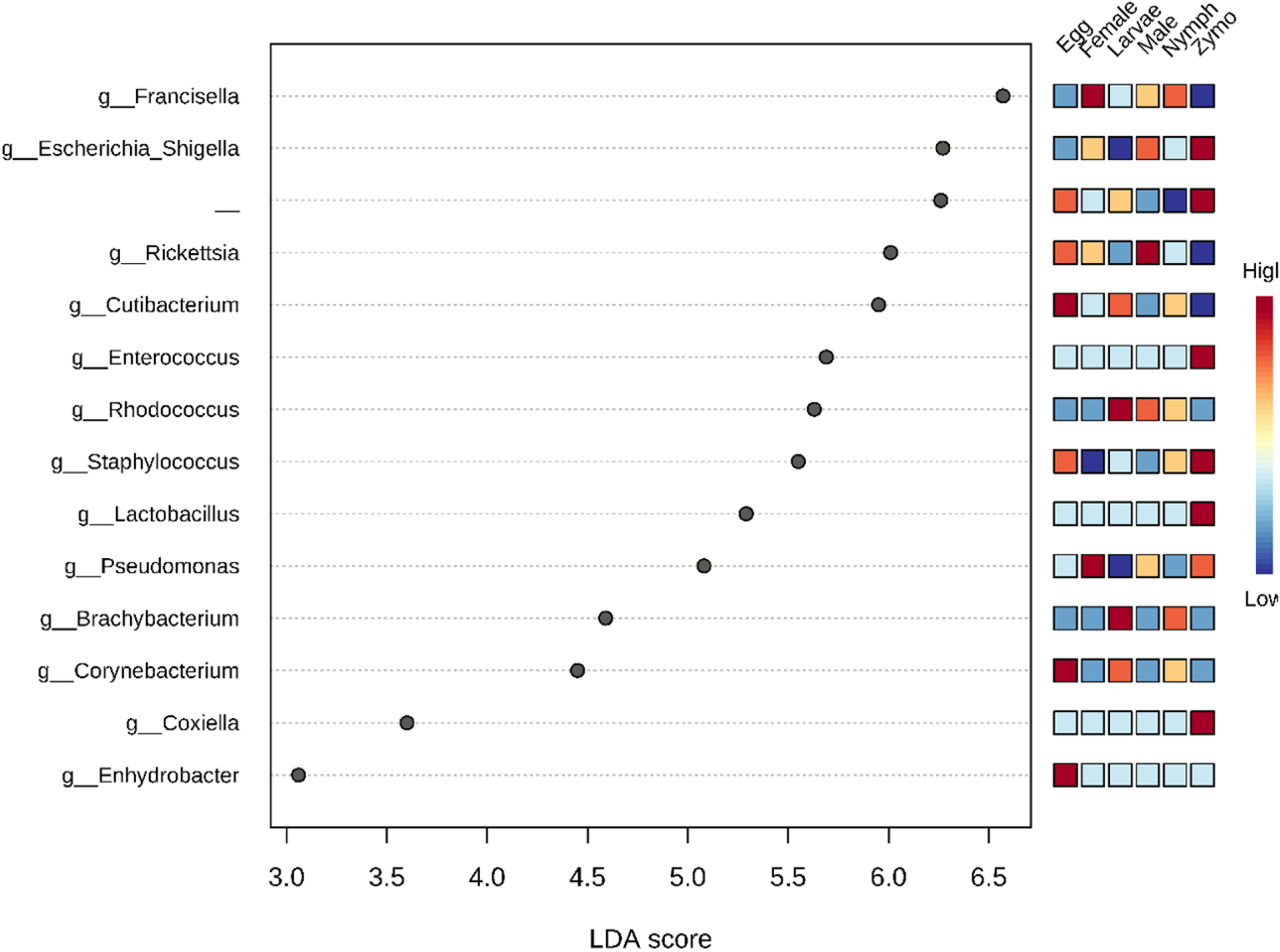
Graphical summary of LEfSe analysis across the developmental stages of *Am. maculatum*. Significant taxa are ranked in decreasing order by their LDA scores (x axis). The mini heat map to the right of the plot indicates whether the number of taxa are higher (red) or lower (blue) in each group.

**Figure S8:**
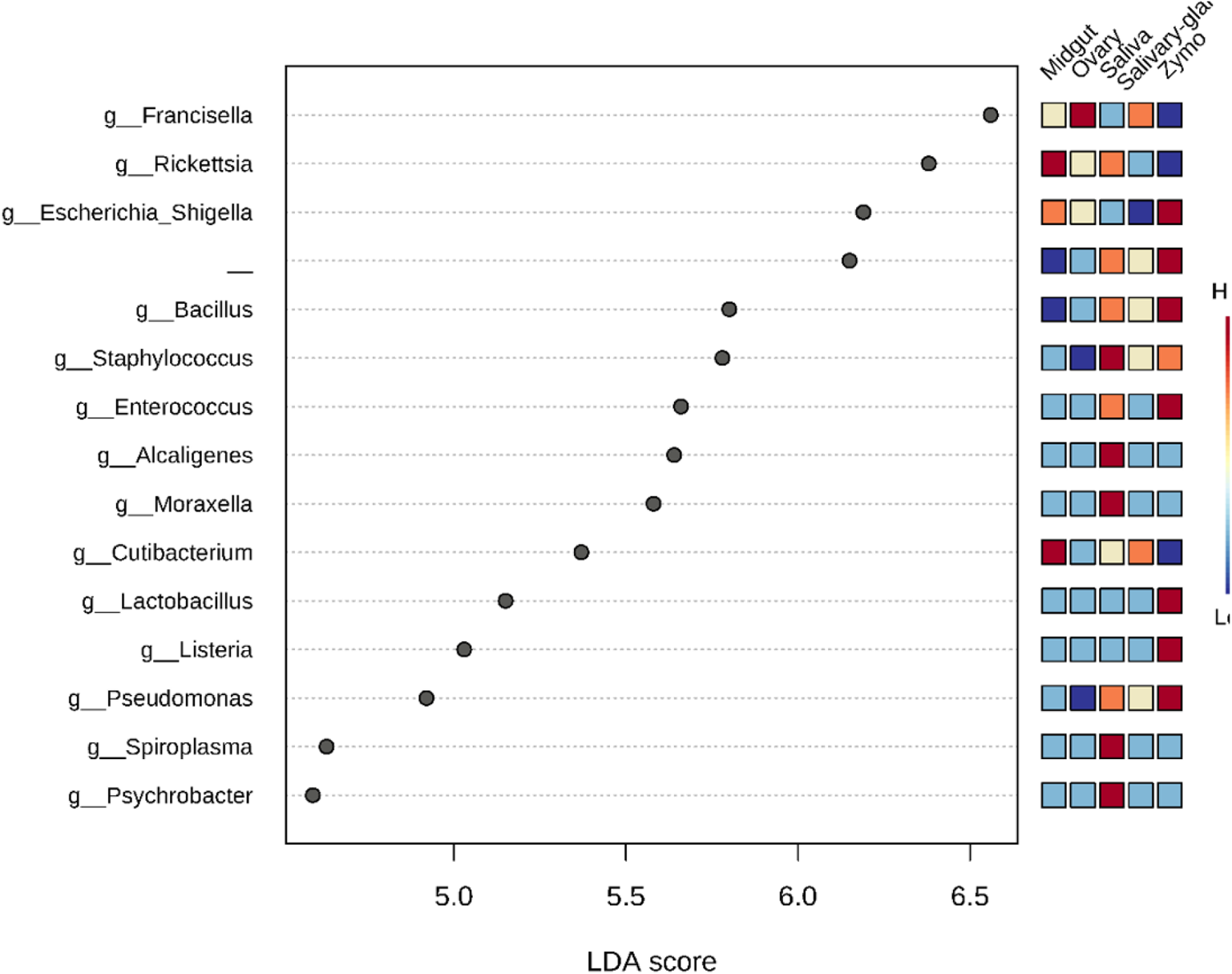
Graphical summary of LEfSe analysis from the dissected tissues of *Am. maculatum*. Significant taxa are ranked in decreasing order by their LDA scores (x axis). The mini heat map to the right of the plot indicates whether the number of taxa are higher (red) or lower (blue) in each group.

**Figure S9:**
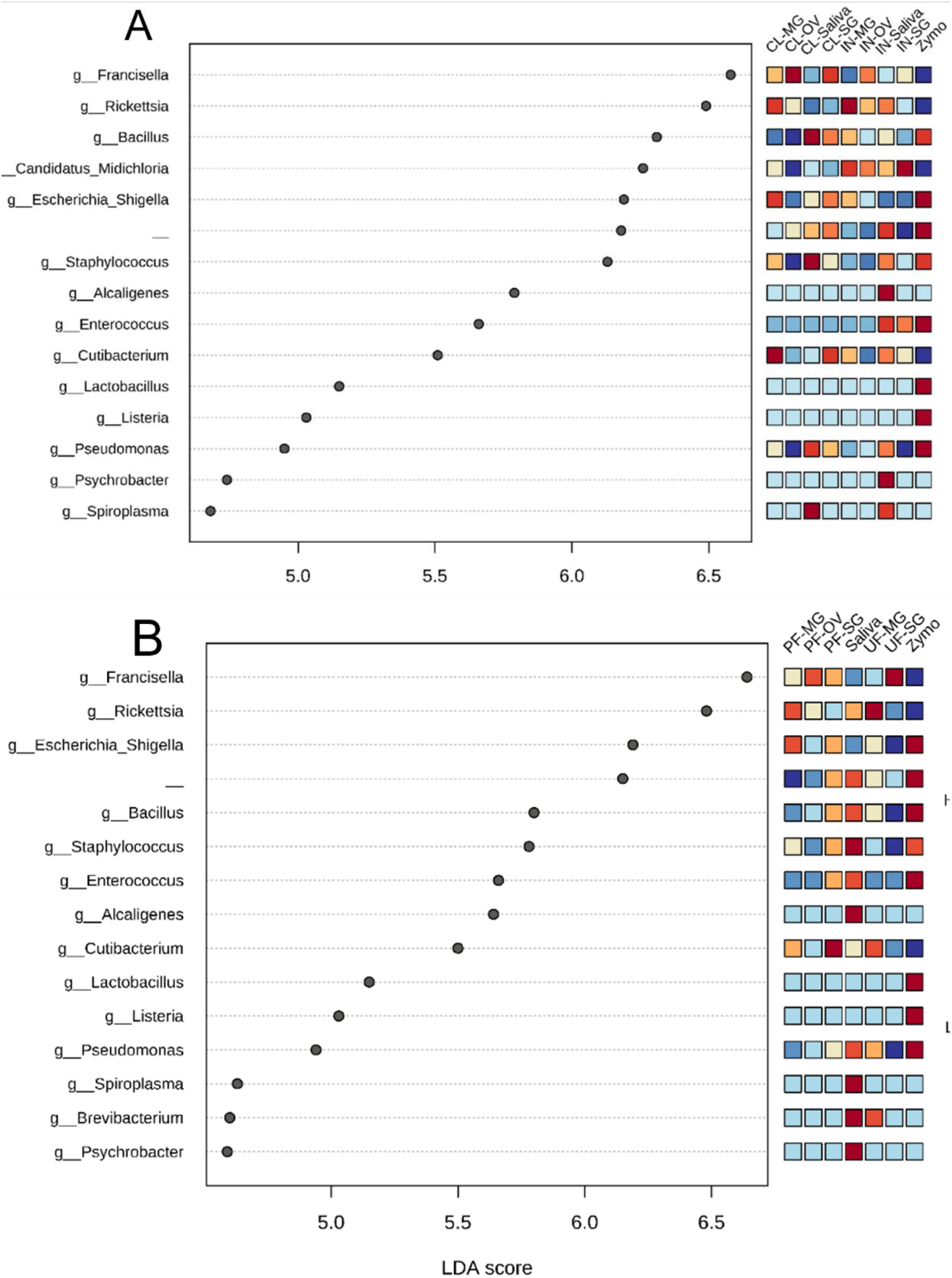
Graphical summary of LEfSe analysis from the dissected tissues of *Am. maculatum*. A) *R. parkeri*-infected and uninfected and B) unfed and partially fed *Am. maculatum*. Significant taxa are ranked in decreasing order by their LDA scores (x axis). The mini heat map to the right of the plot indicates whether the number of taxa are higher (red) or lower (blue) in each group.

**Figure S10:**
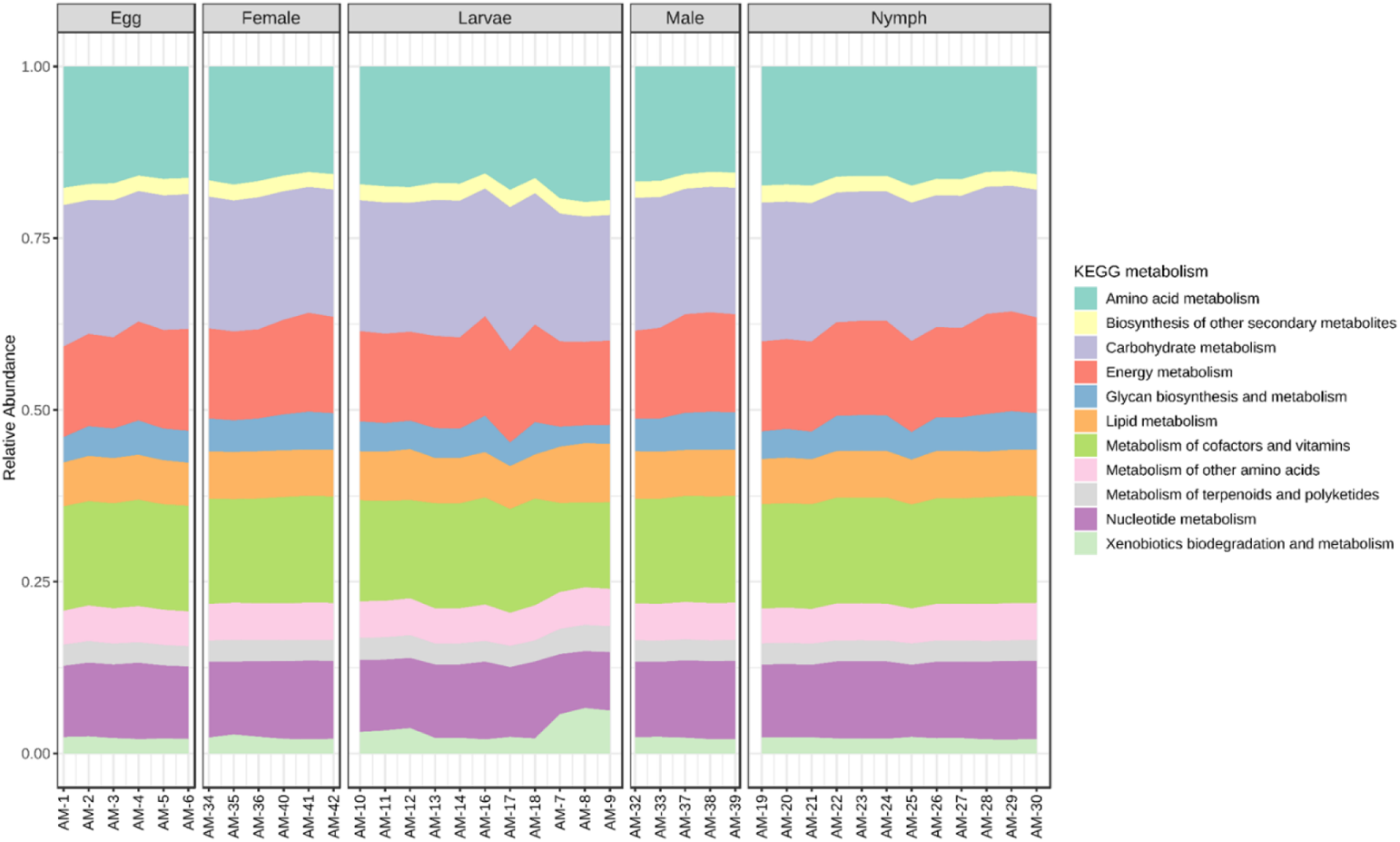
Histogram indicating functional differences of the *Am. maculatum* microbiota across developmental stages. KEGG metabolic categories were obtained from 16S rRNA gene sequences using the PICRUSt2 pipeline.

**Figure S11:**
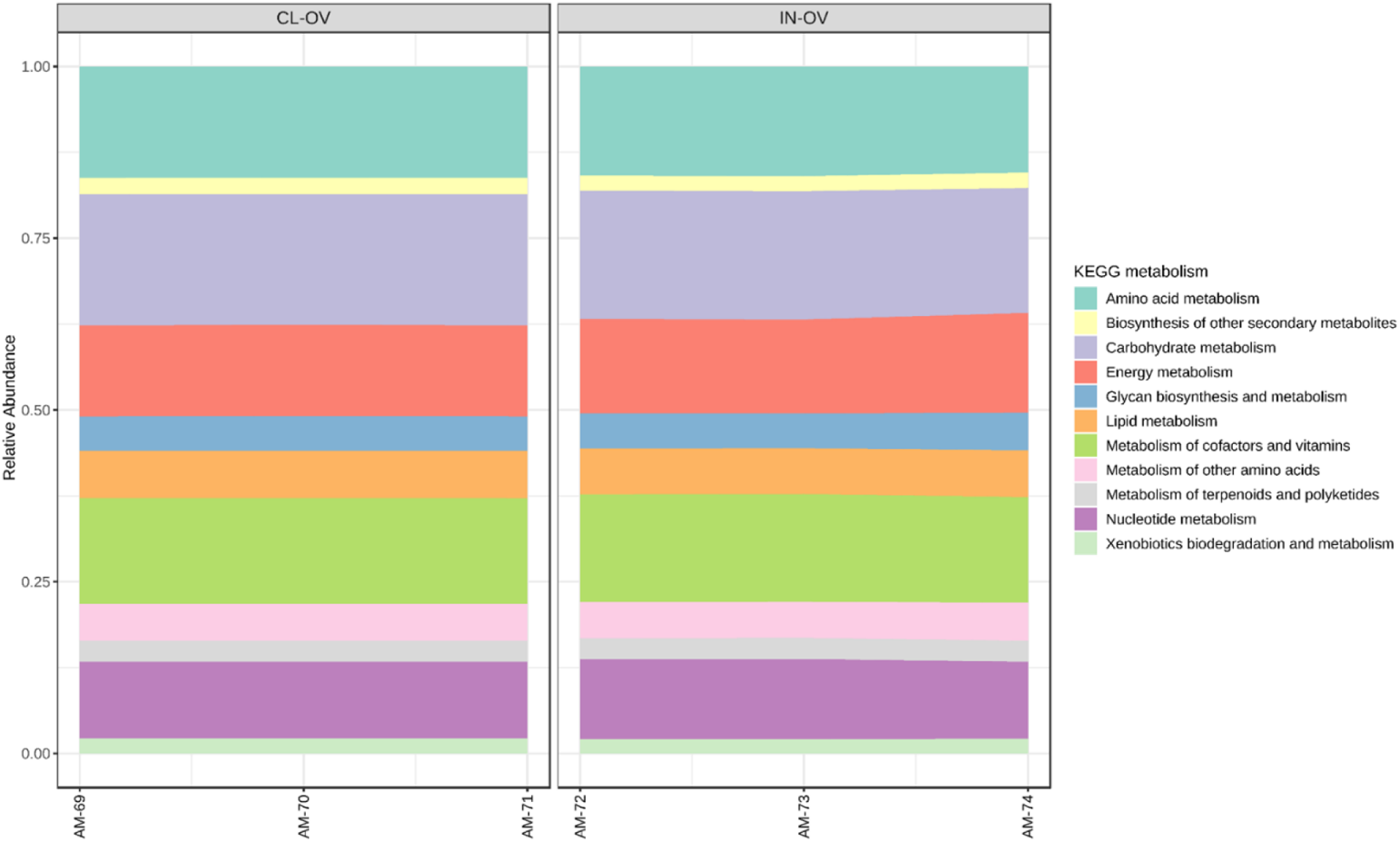
Histogram indicating functional differences in the ovaries of the *Am. maculatum* microbiota with and without *R. parkeri* infection. KEGG metabolic categories were obtained from 16S rRNA gene sequences using the PICRUSt2 pipeline.

## Additional Table

**Table S1:** Demultiplexed sequence counts summary

| Table S1: Demultiplexed sequence counts summary |  |  |
| --- | --- | --- |
|  | Forward reads | Reverse reads |
| Mean | 45453.5 | 45453.5 |
| Maximum | 112242 | 112242 |
| Total | 4090811 | 4090811 |

## Notes

### Competing Interest Statement

The authors have declared no competing interest.

